# TKI-Tolerant Persisters Emerge from a PKCα-Dependent and Highly Plastic Subpopulation of Stem-Like Cells in NSCLC

**DOI:** 10.64898/2026.05.20.726497

**Authors:** Mojtaba Sadeghi, Mohamed Salama, Sagarika Choudhury, Amy Huang, Jie Yang, Yusuf A. Hannun

## Abstract

Reversible drug-tolerant persister states are emerging as key drivers of limited therapeutic durability, offering a complementary non-genetic perspective distinct from traditional models of acquired resistance. This is of particular interest in lung adenocarcinoma where EGFR tyrosine kinase inhibitors (TKIs) elicit dramatic responses, yet residual surviving cells persist and ultimately seed relapse. To define mechanisms that enable survival during this earliest residual-disease phase, we focused on the drug-tolerant persister population that remains after EGFR TKI exposure and can later give rise to outgrowth. Initial observations of elevated transcript levels of *PRKCA*, which encodes PKCα, in established TKI-resistant models, together with markedly delayed tumor relapse following PKCα suppression *in vivo*, nominated PKCα as a candidate regulator of the persister-to-relapse transition. Genetic ablation of *PRKCA* or its inhibition with enzastaurin reduced residual survival and outgrowth after TKI exposure, indicating that PKCα functions as an early dependency of drug-tolerant persisters rather than as a general mediator of acquired resistance. Mechanistically, PKCα was required for persister-associated EMT, migratory capacity, and robust induction of ALDH1A1, the latter constraining oxidative stress and enhancing persister survival. Functionally, PKCα was specifically necessary for survival of a rare, pre-existing CD44^High^ stem-like subpopulation that exhibited marked plasticity and ultimately seeded persistence. Together, these data identify a PKCα-dependent EMT/stemness/ROS pathway as a critical survival program in EGFR TKI-tolerant persister cells and support therapeutic strategies aimed at eliminating residual disease to prolong clinical responses.

## 1 Introduction

Tyrosine kinase inhibitors (TKIs) such as erlotinib and osimertinib have proven highly effective in treating non-small cell lung cancer (NSCLC) driven by specific mutations in the epidermal growth factor receptor (*EGFR*).^1–3^ However, despite impressive initial responses, patients inevitably relapse.^4–6^ While recurrence is often attributed to the presence or evolution of resistant clones, recent findings support a role for cancer persister cells, which are quiescent, drug-tolerant cells that can rebound once treatment is reduced or halted.^7,8^ Although the mechanisms by which persisters arise in response to TKIs are multifactorial and poorly understood, it is widely accepted that reversible, non-mutational mechanisms underlie their emergence.^9^ These changes may promote drug tolerance by enhancing stress-response programs, including autophagy, epithelial-to-mesenchymal transition (EMT), stemness, and alterations in cell-signaling pathways that support survival.^10,11^ Overcoming persistence is crucial for improving the long-term efficacy of TKIs, and recent research has begun to explore therapeutic strategies that target drug-tolerant cells by defining pathways that sustain persister survival.^12,13^

A key step toward improving the durability of TKI responses is to identify the early, targetable dependencies that enable drug-tolerant persisters to emerge before stable resistance is established. Here, building on our prior work showing that mutant EGFR preferentially engages PLCγ and PKCα due to receptor-intrinsic biased signaling,^14,15^ we identify PKCα not as a general resistance factor, as previously reported,^16–21^ but as a critical mediator required during persister genesis, where its loss or inhibition reduces residual survival, suppresses clonogenic persister outgrowth, and delays tumor relapse. Our findings suggest that this requirement is concentrated in a rare, pre-existing CD44^High^ stem-like subpopulation that is intrinsically more tolerant to TKI treatment and uniquely capable of drug-induced plasticity. In this subpopulation, PKCα is necessary for a TKI-induced EMT, migratory functions, and induction of *ALDH1A1* in a program that buffers oxidative stress and allows cells to survive EGFR inhibition. Together, these findings provide direct preclinical evidence that targeting a persister-enabling dependency can reduce residual disease and delay relapse.

## 2 Results

### 2.1 PKCα loss delays *in vivo* relapse but does not reverse established resistance

To test the specific contribution of PKCα to the emergence of resistance *in vivo*, HCC827 cells expressing an empty vector (EV) or a guide directed against *PRKCA* (encoding PKCα) were subcutaneously implanted in athymic mice, followed by daily treatment with erlotinib or osimertinib. Loss of *PRKCA* drastically delayed relapse of tumors, but *PRKCA* knockout (KO) alone had minimal effect on tumor growth, suggesting that PKCα is not an evident standalone dependency in these models but plays an important role in mediating the efficacy and duration of the response to TKIs (Figures 1A and S1A). Similarly, erlotinib or osimertinib co-treatment with enzastaurin, a clinically evaluated PKC inhibitor, delayed breakthrough of TKI-treated HCC827 xenografts with a complete response observed in 2 of 7 cases (Figures 1B, S1B, and S1C). No body-weight loss was observed, and blood chemistry in the ERL/ERL + ENZ cohort did not reveal a broad toxicity signal (Figures S1D-G).^22^ As expected, enzastaurin alone did not demonstrate significant anti-tumor activity, in line with loss of PKCα and previous reports on the inefficacy of the inhibitor as a single-agent anticancer therapeutic.^23–26^ Furthermore, secondary mutations conferring resistance to erlotinib and osimertinib (T790M and C797X, respectively)^27,28^ were not detected in any of the relapsing tumors, suggesting that although loss or inhibition of PKCα drastically delayed relapse, this effect cannot be attributed to the selection of intrinsically resistant clones (Table S1).

**Figure 1.**
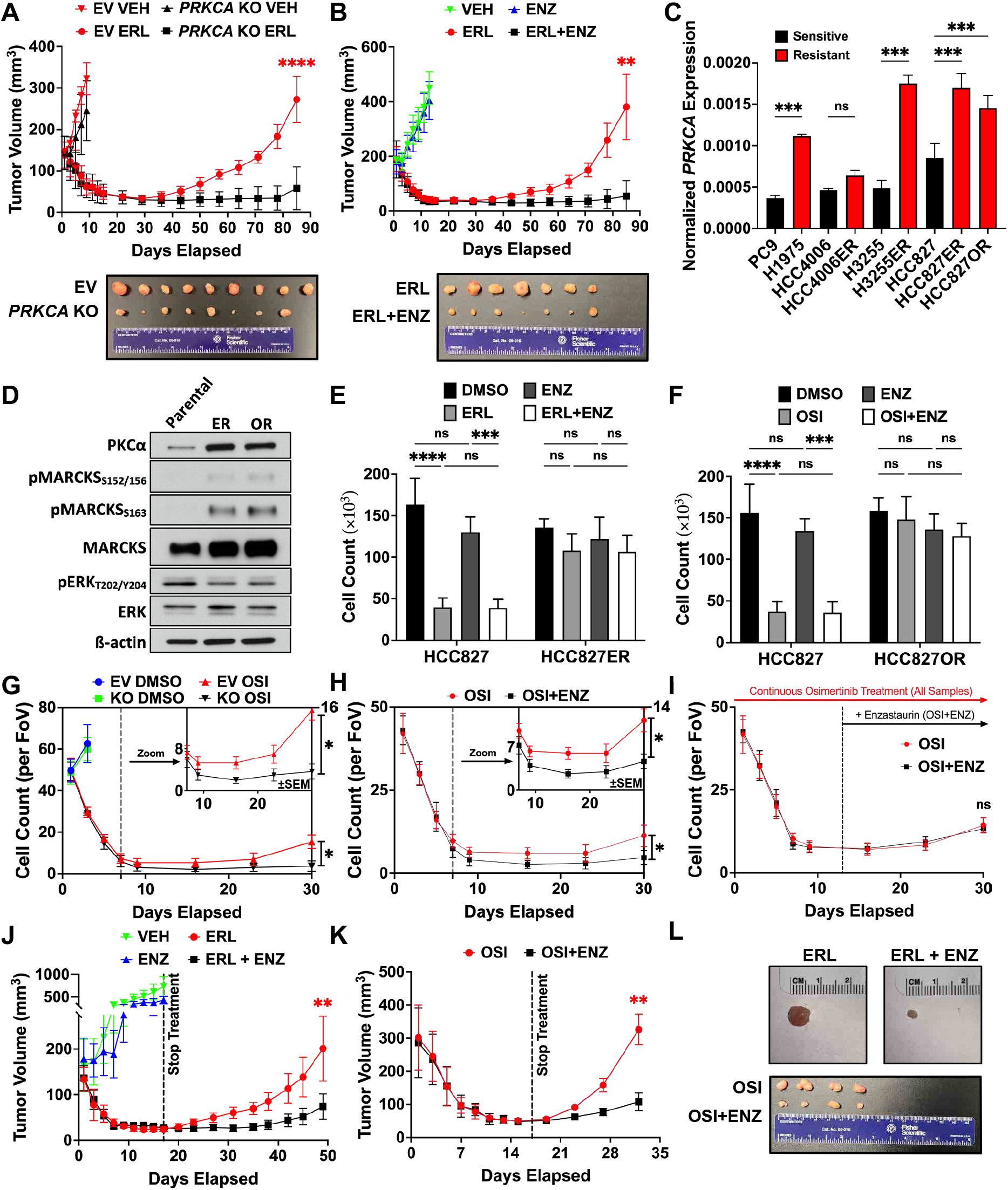
PKCα is upregulated in TKI-resistant cells and promotes tumor relapse. **(A)** Tumor volumes of HCC827 empty vector (EV) and *PRKCA* knockout (KO) xenograAs treated with vehicle (VEH; 10% acacia) or erlotinib (ERL; 40 mg/kg/day) for 85 days (*n* = 9). **(B)** Tumor volumes of HCC827 xenograAs treated with VEH (10% acacia), ERL (40 mg/kg/day), enzastaurin (ENZ; 125 mg/kg twice daily), or ERL + ENZ for 85 days (*n* = 7). **(C)** Mean normalized expression of *PRKCA* in mutant EGFR (mEGFR)-positive cell lines that are TKI-sensitive (black) or TKI-resistant (red). Culture-generated erlotinib-resistant (ER) and osimertinib-resistant (OR) cell lines are indicated (*n* = 3). **(D)** Immunoblot of parental HCC827 and TKI-resistant HCC827ER and HCC827OR cells assessing PKCα expression and downstream signaling (*n* = 2). **(E)** Cell counts of HCC827 and HCC827ER cells treated with dimethyl sulfoxide (DMSO), ERL (100 nM), ENZ (1 µM), or ERL + ENZ for 48 h (*n* = 3). **(F)** Cell counts of HCC827 and HCC827OR cells treated with DMSO, osimertinib (OSI; 10 nM), ENZ (1 µM), or OSI + ENZ for 48 h (*n* = 3). **(G)** Cell counts of HCC827 EV or *PRKCA* KO cells during continuous OSI treatment (10 nM) for 30 days. Mean adherent cells per field of view (FOV) were quantified from three random FOVs at 10× magnification at the indicated Ime points (*n* = 3). **(H)** Cell counts of HCC827 cells treated with OSI (10 nM) alone or OSI + ENZ (1 µM) for 30 days, quantified as in G (*n* = 3). **(I)** Cell counts as in H, with ENZ initiated on day 13 in the OSI + ENZ group (*n* = 3). **(J)** Tumor volumes of HCC827 xenograAs treated with VEH (10% acacia), ERL (40 mg/kg/day), ENZ (125 mg/kg twice daily), or ERL + ENZ; treatments were withdrawn aAer day 13 (*n* = 8). **(K)** Tumor volumes of TM00193 patient-derived xenograAs (EGFR_del746-750_; treatment-naïve) treated with OSI (25 mg/kg/day) or OSI + ENZ (125 mg/kg twice daily); treatments were withdrawn aAer day 17 (*n* = 4). **(L)** Representative images of tumors from J and K. Data are presented as mean ± SD (or ± SEM where indicated). *p-va*lues for A,B,G–K were calculated using mixed-effects models. *p-va*lues for C were calculated using multiple t-tests comparing resistant cell lines with their matched parental/sensitive counterparts. *p-va*lues for E and F were determined by two-way ANOVA with Šidák’s correction. \**p* < 0.05, \*\**p* < 0.01, \*\*\**p* < 0.001, \*\*\*\**p* < 0.0001, ^ns^*p* > 0.05 or not significant.

Sections obtained from recurrent tumors demonstrated increased PKCα in osimertinib-relapsed samples as compared with vehicle-treated or enzastaurin co-treated tumors, suggesting a potential role in resistance (Figure S1H). In line with this finding, publicly available RNA-seq data from the Cancer Cell Line Encyclopedia (CCLE) revealed elevated *PRKCA* transcript in TKI-resistant NSCLC cell lines (Figure S2A). To corroborate these results in culture and eliminate the possibility of confounding effects from cross-cell-line comparisons, erlotinib and/or osimertinib-resistant cell lines (denoted as ER and OR, respectively) were established from the parental HCC827, HCC4006, and H3255 cells through continuous treatment with 100 nM erlotinib or 10 nM osimertinib (doses determined to attenuate EGFR autophosphorylation) over the course of two months (Figures S2B and S2C). The resulting resistant cell lines demonstrated at least a 40-fold increase in IC_50_ when treated with erlotinib or osimertinib (Figures S2D, S2E, S2F, and S2G). PKCα expression analysis by real-time RT-qPCR revealed a clear upregulation of *PRKCA* transcript levels in both *de novo* and culture-generated TKI-resistant cell lines (Figure 1C). Of note, PKCα retained activation status despite erlotinib treatment in resistant cells, as assessed by visualization of plasma membrane translocation, where the enzyme interacts with activators and phosphorylates substrates (Figure S2H). This is in stark contrast to non-resistant cells, where the activity of PKCα is strictly dependent on active mEGFR. We thus asked whether PKCα provides an escape mechanism that modulates sensitivity to TKIs. PKCα is robustly overexpressed more than 10-fold in HCC827ER and HCC827OR cells, with enhanced activation of PKCα substrate MARCKS_S163_, but little to no change in the expression and activity of ERK (Figure 1D). However, enzastaurin treatment did not resensitize HCC827ER, HCC827OR, or HCC4006ER cells to TKIs (Figures 1E, 1F, and S2I). Moreover, addition of enzastaurin did not enhance the acute response of sensitive cells to TKIs, thus arguing against appreciable synergy between the inhibitors. While these findings make a case against PKCα as a modulator of TKI resistance, the robust elevation of PKCα in resistant cell lines and the dramatic *in vivo* responses with suppression of PKCα in TKI-treated xenografts raised the question as to the specific role of PKCα during the emergence of resistance.

### 2.2 PKCα suppression limits residual cells following TKI treatment

To study the specific functional role of PKCα in the response to chronic TKI treatment, the effects of loss or inhibition of PKCα on the response to osimertinib were assessed. Regardless of inhibition or deletion of PKCα, a similar initial response was observed during chronic treatment of HCC827 cells, with most cells dying rapidly upon initiation of osimertinib and until day 9, when the cell counts plateaued (Figures 1G and 1H). However, a small but non-negligible population of surviving cells remained; this population was markedly reduced by loss or inhibition of PKCα. While this reached statistical significance only at the 30-day time point, an appreciable difference can be noted as early as day 9 (Figures 1G and 1H insets). When enzastaurin treatment was delayed until 13 days after TKI treatment initiation, the observed difference in residual cells was no longer observed, thus strongly suggesting that PKCα inhibition is critical upfront, within the initial exposure to osimertinib, to limit survival of residual cells (Figure 1I). To evaluate these findings in animal models, xenografts harboring HCC827 cells or patient-derived tumors (PDXs) were treated with erlotinib or osimertinib, respectively, and the effect of co-treatment with enzastaurin on tumor growth was assessed. Despite withholding treatment after 13 days (17 days for PDXs), inhibition of PKCα significantly reduced tumor regrowth in both models (Figures 1J, 1K, and 1L). Since suppression of PKCα in the initial days of TKI therapy was sufficient to reduce the fraction of surviving cells *in vitro* and tumor regrowth *in vivo*, we hypothesized that PKCα may prime treatment-naïve cells for survival, thus regulating the genesis of drug-tolerant persister cells as opposed to modulating resistance once it has been established.

### 2.3 PKCα is critical for the genesis of TKI-tolerant persisters

To specifically assay persistence, a clonogenic protocol was adopted with treatment durations of 14 days followed by a brief recovery period to allow for the formation of colonies from drug-tolerant clones, with each surviving colony at endpoint originating from a single residual clone (Figure 2A). Quantitation of these colonies revealed that the loss of PKCα diminished persistence by 50–70% with either erlotinib or osimertinib treatment (Figures 2B and 2C), and re-expression of PKCα rescued this phenotype (Figure 2D), thus strongly supporting the hypothesis that PKCα is necessary for the survival of residual clones following prolonged EGFR TKI exposure. To test these findings pharmacologically using a clinically evaluated PKC inhibitor, enzastaurin was utilized. While enzastaurin treatment alone demonstrated virtually no effects on clonogenic survival, the inhibitor markedly potentiated the effects of TKIs, reducing clonogenic survival by 70% when cells were treated with erlotinib or osimertinib in combination with enzastaurin (Figures 2E, 2F, and 2G). Collectively, these results strongly suggest that suppression of PKCα effectively reduces the number of viable clones following erlotinib or osimertinib therapy, enhancing the effects of TKIs and limiting the remaining reservoir of cells that are poised to drive tumor recurrence.

**Figure 2.**
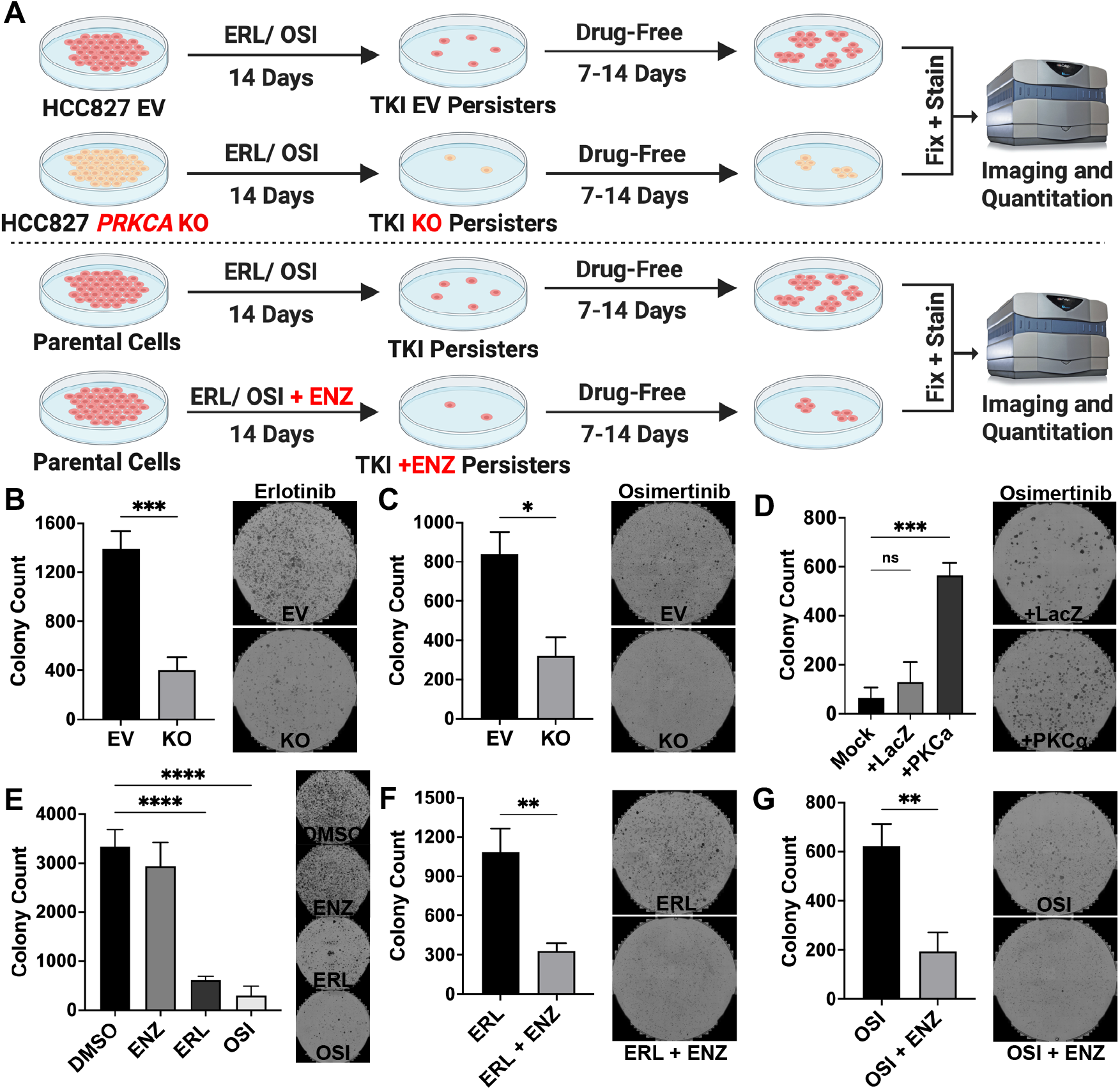
PKCα drives clonogenic survival of TKI-treated cells. **(A)** Schematic of the clonogenic assay used to develop persisters and test persistence under different treatment conditions and/or genetic perturbations. **(B**,**C)** Clonogenic survival of HCC827 EV and *PRKCA* KO cells treated with erlotinib (ERL; 100 nM) **(B)** or osimertinib (OSI; 10 nM) **(C)** for 14 days, followed by 7–14 days of recovery. Representative stained plates are shown (*n* = 3). **(D)** Clonogenic survival of *PRKCA* KO cells expressing LacZ control or re-expressing *PRKCA* following OSI treatment (10 nM). Mock indicates infection control without rescue construct (*n* = 3). **(E)** Clonogenic survival of HCC827 cells treated with DMSO, ENZ (1 µM), ERL (100 nM), or OSI (10 nM) (*n* = 3). **(F**,**G)** Clonogenic survival of HCC827 cells treated with ERL (100 nM) **(F)** or OSI (10 nM) **(G)** alone or in combination with ENZ (1 µM) (*n* = 3). Data are presented as mean ± SD. *p-va*lues for B,C,F, and G were determined by unpaired two-sided t-tests. *p-va*lues for D and E were calculated using one-way ANOVA with Šidák’s correction. \**p* < 0.05, \*\**p* < 0.01, \*\*\**p* < 0.001, \*\*\*\**p* < 0.0001, ^ns^*p* > 0.05.

### 2.4 PKCα is upregulated, active, and decoupled from mEGFR in TKI-tolerant persisters

To investigate the role of PKCα in persistence, we generated monoclonal erlotinib persister clones (EP1-9) and osimertinib persister clones (OP1-9), as well as polyclonal persister populations (denoted EP or OP, respectively) from HCC827 cells (Figure 3A). The appropriate vehicle-generated controls were also prepared through treatment with DMSO. Unlike resistant cells, HCC827 EP and OP regained sensitivity after roughly 15 passages, an expected distinct property of persisters that distinguishes them from resistant cells (Figures S3A and S3B). Moreover, a roughly 30-fold increase in the expression of carnitine palmitoyltransferase 1A (*CPT1A*), known to be highly upregulated in drug-tolerant cells, was observed in persisters, thus further validating the unique identity of these cells (Figures 3B and 3C).^29^ Interestingly, in both erlotinib and osimertinib persisters, an average 4-fold increase in PKCα was observed by Western blot (Figures 3D and 3E), with a 2-to 3-fold increase in *PRKCA* mRNA (Figures 3F and 3G). This increase was specific to PKCα and was not observed for other PKC isozymes (Figure S3C). Notably, acute treatment with erlotinib or osimertinib for 24 hours did not induce upregulation of *PRKCA* (Figure S3D), suggesting that the increase in PKCα is not a direct result of drug exposure and response, but possibly a consequence of selection of cells with high PKCα expression levels. Furthermore, PKCα was predominantly localized to the plasma membrane, and robust phosphorylation of MARCKS_S159/163_ and EGFR_T654_ was observed in persisters compared to vehicle-generated controls (Figures 3H, 3I, and 3J), suggesting that the kinase is not only overexpressed, but also shows higher activity. Despite demonstrating increased expression and activity, inhibition of PKCα in persisters did not elicit a more significant effect on the proliferation of EP and OP cells compared to vehicle-generated control cells and failed to resensitize early-passage persisters to TKIs (Figures 3K, 3L, and S3E). These results strongly suggest that while PKCα is critical for the generation of persisters, it is not necessary for the maintenance of the drug-tolerant state, consistent with the lack of an appreciable role for PKCα in established resistant cells. To investigate whether activity of PKCα in persisters could be attributed to acquired or selected EGFR insensitivity to TKIs, known EGFR phosphosites that couple to the activation of PKCα through PLCγ were assessed. These results indicate that TKIs inhibit mEGFR to a similar degree in persisters as compared with control cells (Figures 3M and S3F). Moreover, and unlike the case for the control or naïve cells where localization of PKCα to the plasma membrane and signaling to p70S6K are dependent on mEGFR, PKCα retained membrane localization (Figures 3N and S3G) and p70S6K_T389_ was phosphorylated in persisters despite TKI treatment (Figure 3M), and this phosphorylation was inhibited by enzastaurin (Figure S3H). These results demonstrate that the activation of PKCα in persister cells is driven by mEGFR-independent mechanisms but is still necessary for critical downstream signaling.

**Figure 3.**
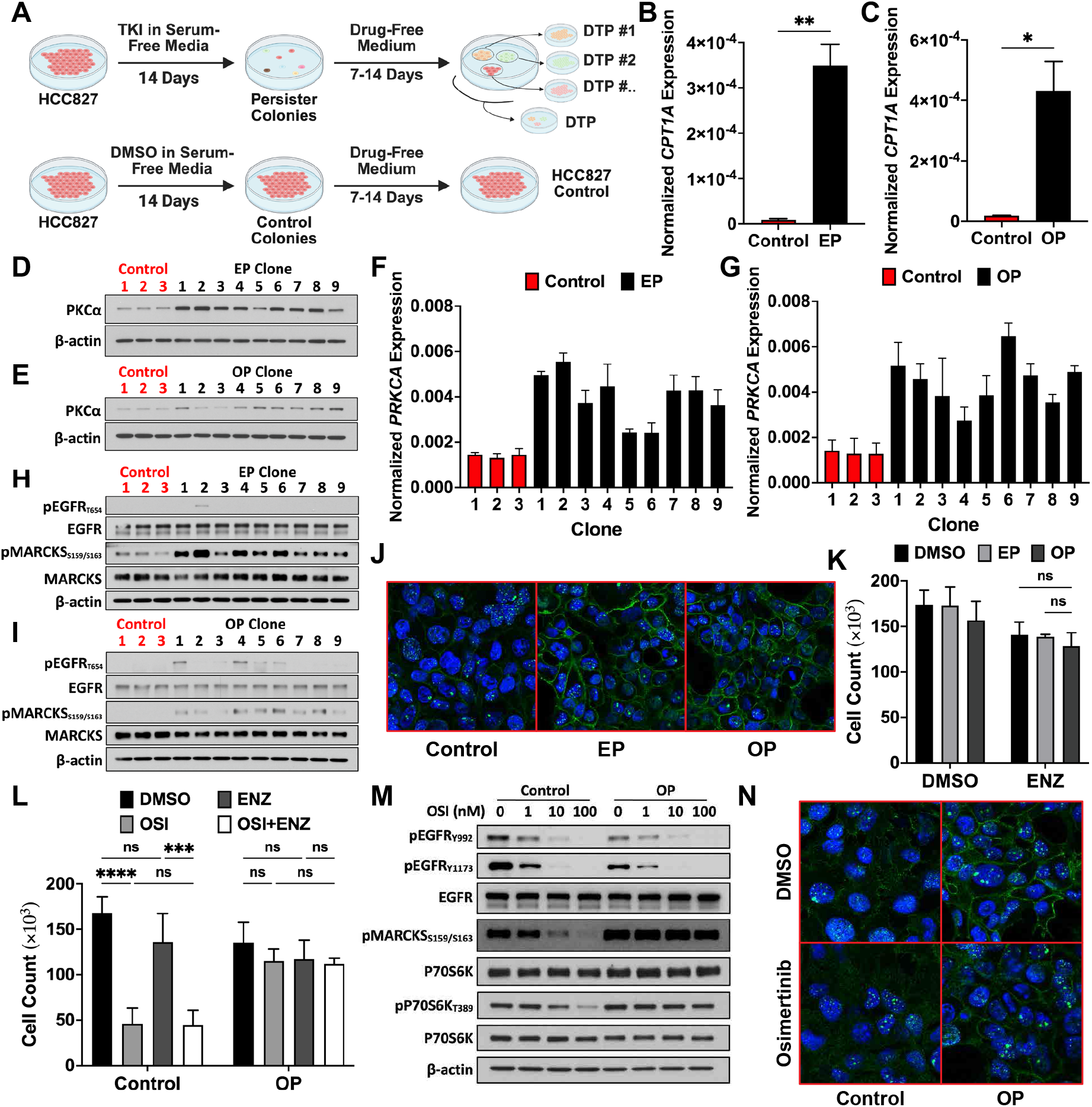
PKCα is highly expressed and acMve independent of mEGFR in persisters. **(A)** A schematic showing the generation of drug-tolerant persisters (DTPs) using EGFR TKIs ERL or OSI. Monoclonal ERL or OSI persisters are denoted EP1–9 or OP1–9, respectively. Polyclonal persisters are denoted EP and OP. **(B**,**C)** Normalized expression (to *ACTB*) of *CPT1A* in EP **(B)** or OP **(C)** cells (*n* = 3). **(D**,**E)** PKCα protein levels in EP **(D)** and OP **(E)** persister clones (*n* = 3). **(F**,**G)** Normalized *PRKCA* expression in monoclonal EP **(F)** and OP **(G)** persisters (*n* = 3). **(H**,**I)** Immunoblot depicting phosphorylation of MARCKS_S159/S163_ and EGFR_T654_ (PKC site) in control cells and EP clones **(H)** or OP clones **(I)** (*n* = 2). **(J)** Immunofluorescence staining of PKCα (green) in HCC827 control, EP, and OP cells. Nuclei are stained with DAPI (*n* = 3). **(K)** Cell counts of HCC827 control, EP, or OP cells following treatment with DMSO or ENZ (1 µM) for 48 h (*n* = 3). **(L)** Cell counts of HCC827 control and OP cells treated with DMSO, OSI (10 nM), ENZ (1 µM), or OSI + ENZ (*n* = 3). **(M)** Immunoblot showing phosphorylation of EGFR at two sites that couple to PKC signaling and downstream effectors of PKC in control and OP cells following OSI treatment at varying doses (*n* = 2). **(N)** PKCα membrane localization (green) in HCC827 control and OP cells following OSI treatment (10 nM) (*n* = 3). Data are presented as mean ± SD. *p-va*lues for B and C were calculated using unpaired two-sided t-tests. *p-va*lues for K and L were determined by two-way ANOVA with Šidák’s correction. \**p* < 0.05, \*\**p* < 0.01, \*\*\**p* < 0.001, \*\*\*\**p* < 0.0001, ^ns^*p* > 0.05.

### 2.5 Persisters demonstrate PKCα-dependent EMT

In agreement with previous reports regarding persistence, nearly all persister clones exhibited mesenchymal morphology with a clear spindle-shaped structure and grew as isolated cells, whereas the control cells retained their regular epithelial-like form and grew in patches (Figures 4A and S4A). Furthermore, microarray analysis of control and persister cell lines revealed a significant enrichment of EMT-associated gene signatures including cell motility, migration, and differentiation (Figure S4B). In line with these results, HCC827 EP and OP clones uniformly expressed higher vimentin and showed undetectable levels of E-cadherin when compared to the vehicle-generated controls, suggesting acquisition of a mesenchymal state following TKI treatment (Figures 4B and 4C). Importantly, persisters exhibited significant mesenchymal characteristics, a phenotype that was not induced in *PRKCA* KO cells (Figure S4C). This, along with previous reports on PKCα as a promoter of EMT and a driver of migration in breast cancer, ^19,30^ led us to ask if the kinase plays a similar role in the current model of persistence.

**Figure 4.**
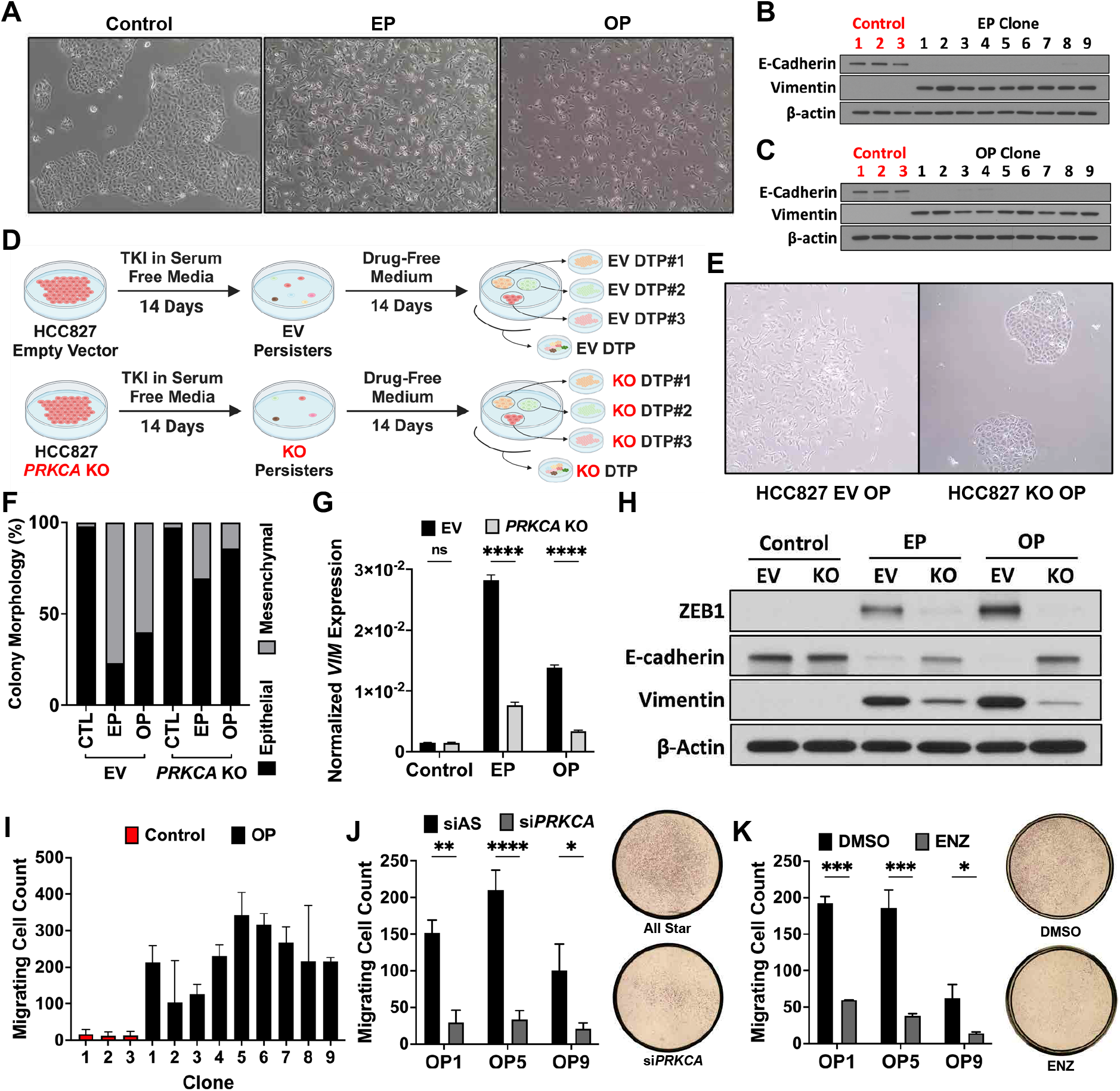
Persisters undergo PKCα-dependent EMT and migraMon. **(A)** Brighsield images of HCC827 control, ERL persisters (EP), and OSI persisters (OP). **(B**,**C)** Western blot analysis of E-cadherin and vimentin in control vs erlotinib persister clones (EP1-9) **(B)** or control vs osimertinib persister clones (OP1-9) **(C)** cells (*n* = 2). **(D)** Schematic depicting generation of persisters from HCC827 EV and *PRKCA* KO cells; *PRKCA* loss was induced prior to persister generation. **(E)** Brighsield images of EV and *PRKCA* KO OSI persisters. **(F)** Quantification of EV and *PRKCA* KO persisters with epithelial or mesenchymal morphology. States were assigned by measuring intercellular distances within colonies using ImageJ; predominantly non-zero distances were scored as mesenchymal (*n* = 10-32). **(G)** Normalized expression (to *ACTB*) of vimentin (*VIM)* mRNA in EV and *PRKCA* KO persisters (*n* = 3). **(H)** Western blot analysis of ZEB1, E-cadherin, and vimentin in EV and *PRKCA* KO persisters (*n* = 3). **(I)** Transwell migration of HCC827 control and OP clones. Cells were allowed 16 h to migrate to the lower chamber and were quantified aAer crystal violet staining and ImageJ analysis (*n* = 3). **(J)** Migrating cell counts of OP clones pretreated with All-Star negative control siRNA (siAS; 20 nM) or *PRKCA* siRNA (*n* = 3). **(K)** Migrating cell counts of OP clones pretreated with DMSO or enzastaurin (ENZ; 1 µM) for 6 h (*n* = 3). Cells were then allowed 16 h to migrate in the continued presence of DMSO or ENZ. Representative stitched images of whole inserts are shown (*n* = 3). Data are presented as mean ± SD. *p-va*lues for G,J, and K were determined by multiple unpaired two-sided t-tests. \**p* < 0.05, \*\**p* < 0.01, \*\*\**p* < 0.001, \*\*\*\**p* < 0.0001, ^ns^*p* > 0.05.

To test whether PKCα contributes to the observed mesenchymal phenotype of persister cells, TKI-tolerant cells were generated from HCC827 EV and *PRKCA* KO cells, where loss of PKCα preceded the generation of persisters (Figure 4D). While EV cells demonstrated robust mesenchymal phenotypes following chronic treatment with TKI, the persister cells generated from *PRKCA* KO maintained their epithelial morphology (Figures 4E and 4F). Furthermore, persisting clones generated from *PRKCA* KO cells expressed relatively lower or undetectable levels of vimentin and ZEB1, and retained expression of E-cadherin, thereby suggesting that PKCα is necessary for TKI-induced EMT (Figures 4G, 4H, S4D, S4E, S4F, and S4G). To determine whether PKCα is also necessary for the maintenance of the mesenchymal state, CRISPR-mediated KO and siRNA-mediated knockdown (KD) of *PRKCA* were performed in HCC827 EP and OP cells (established persisters) (Figures S4H and S4I). Unlike loss of PKCα prior to genesis of persistence, deletion of *PRKCA* in already established persisters only partially reverted the mesenchymal phenotype of HCC827 EP (Figure S4J). Furthermore, downregulation of *PRKCA* in erlotinib or osimertinib persisters did not reduce vimentin or restore E-cadherin expression (Figures S4K and S4L). This suggests that while PKCα is necessary for the induction of EMT-positive persister cells, the expression of the kinase is not required for maintenance of the mesenchymal state. Since cell migration and motility are key manifestations of EMT,^31^ and most persister clones demonstrated a significant ability to migrate (Figures 4I and S4M), we assessed whether PKCα is necessary for this function in persisters. To this end, the expression or activity of the isozyme was suppressed in 3 persister clones with the highest levels of PKCα. Here, KD or inhibition of PKCα significantly reduced the capacity of persisters to migrate (Figures 4J, 4K, S4N, and S4O). While dispensable in the maintenance of at least some mesenchymal markers, the results here suggest that expression and activity of PKCα are required for the migratory capacity of persisters, even though the cells retained a mesenchymal phenotype.

### 2.6 PKCα-mediated induction of ALDH1A1 constrains ROS to promote drug tolerance

Persister cells are thought to leverage an array of signaling networks to survive drug pressure and seed recurrence.^32^ To define the specific involvement of PKCα in these networks, a selective expression screen was performed to test the effects of *PRKCA* KO on the induction of key known modulators of persistence. This screen identified ALDH1A1 (Figure 5A),^33–35^ a marker of cancer stem cells (CSCs) and a functional contributor to drug tolerance in several cancers, as a PKCα-dependent persistence-associated factor. *ALDH1A1* demonstrated over a 50-fold increase in expression within TKI-induced cells. Importantly, loss of PKCα in HCC827 prior to the induction of persistence nearly eliminated the increase in expression of *ALDH1A1* at the protein and mRNA levels within EP and OP cells, and in most monoclonal persisters (Figures 5B, 5C, 5D, S5A, and S5B). Moreover, loss of PKCα or enzastaurin treatment abolished the erlotinib- or osimertinib-induced *ALDH1A1* expression in tumors from both HCC827 xenografts and in PDXs, respectively (Figures 5E and 5F). Collectively, these results support the conclusion that persistence-induced *ALDH1A1* expression requires PKCα.

**Figure 5.**
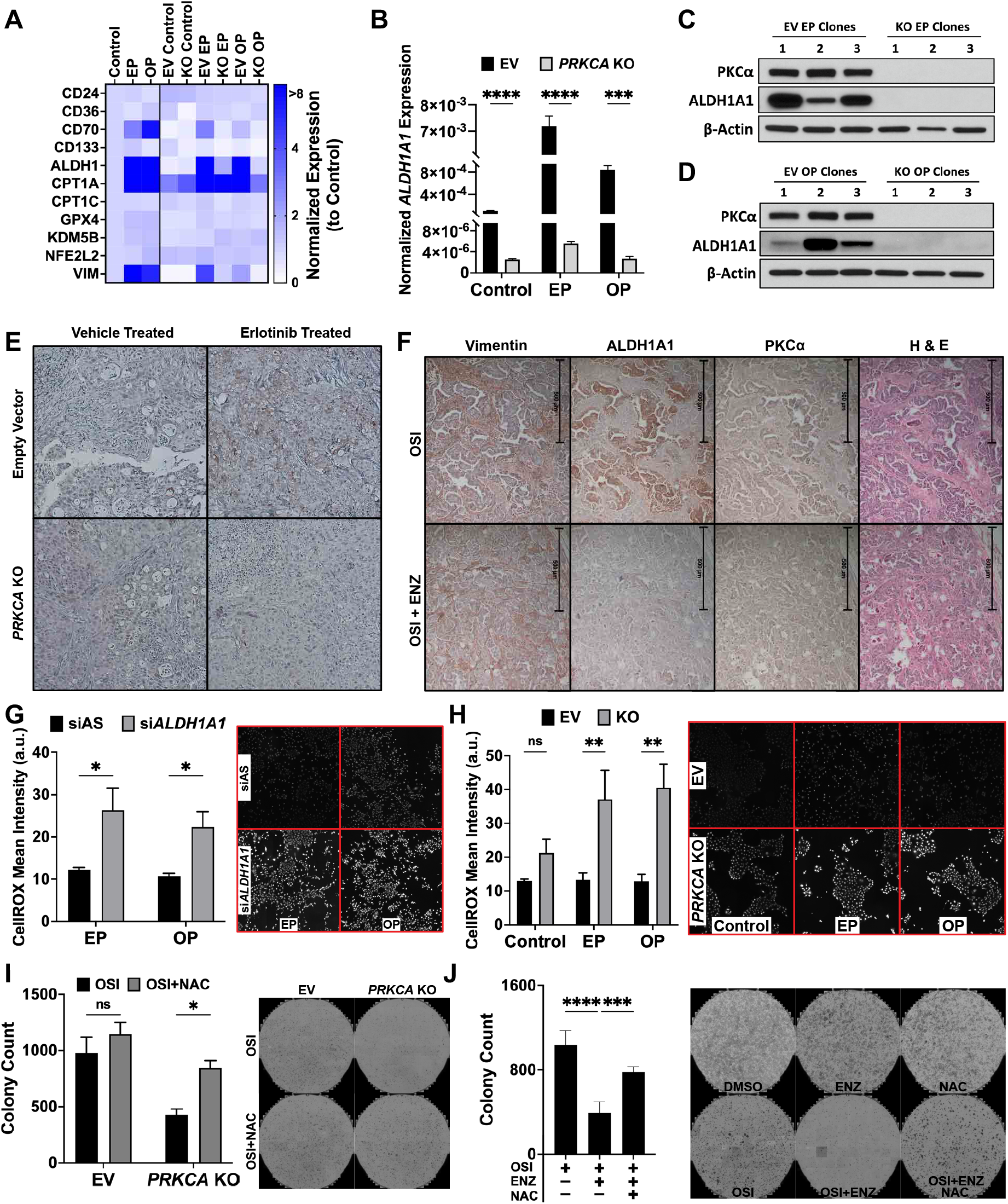
PKCα-dependent ALDH1A1 expression promotes persistence through ROS buffering. **(A)** Selective TaqMan array assessing expression of persistence-associated markers in control, erlotinib persisters (EP), or osimertinib persisters (OP) generated from HCC827 cells expressing an empty vector (EV) or with loss of *PRKCA* (KO). Expression of indicated genes was normalized to *ACTB* and then to the control condition to yield fold change relative to the control sample (*n* = 3). **(B)** Normalized *ALDH1A1* expression (to *ACTB*) in persisters generated from HCC827 EV or *PRKCA* KO cells (*n* = 3). **(C**,**D)** Western blot analysis of PKCα and ALDH1A1 in EV and *PRKCA* KO ERL persister clones **(C)** or OSI persister clones **(D)** (*n* = 3). **(E)** ALDH1A1 staining in HCC827 EV and *PRKCA* KO xenograAs from Figure 1A treated with vehicle or ERL as previously described. Representative sections are shown (*n* = 3). **(F)** Sections from patient-derived xenograAs in Figure 1K treated with OSI alone or OSI + ENZ, stained for indicated targets or with H&E (*n* = 3). **(G)** Mean fluorescence intensity of CellROX Green in EP and OP cells transfected with siAS or *ALDH1A1* siRNA for 48 h, followed by menadione treatment (200 µM) to induce ROS (*n* = 3). **(H)** Mean fluorescence intensity of CellROX Green in Control, EP, and OP cells generated from HCC827 EV or *PRKCA* KO backgrounds (*n* = 3). **(I)** Clonogenic survival of HCC827 EV and *PRKCA* KO cells treated with OSI (10 nM) alone or OSI + N-Acetyl-L-cysteine (NAC; 1 mM) (*n* = 3). **(J)** Clonogenic survival of HCC827 cells treated with OSI (10 nM), OSI + ENZ (1 µM), or OSI + ENZ + NAC (1 mM). Representative plates also include DMSO, ENZ, and NAC control conditions. Data are presented as mean ± SD. *p-va*lues for B and G were calculated using multiple unpaired two-sided t-tests. *p-va*lues for H and I were determined by two-way ANOVA with Šidák’s correction. *p-va*lues for J were calculated using one-way ANOVA with Šidák’s correction. \**p* < 0.05, \*\**p* < 0.01, \*\*\**p* < 0.001, \*\*\*\**p* < 0.0001, ^ns^*p* > 0.05.

Functionally, ALDH1A1 detoxifies reactive lipid-peroxidation-derived aldehydes into less reactive acids, interrupting aldehyde-driven ROS amplification. Therefore, we hypothesized that the robust PKCα-mediated upregulation of *ALDH1A1* upon induction of persisters may support drug tolerance by dampening ROS accumulation. Consistent with this hypothesis, ALDH1A1 knockdown increased CellROX signal in menadione-treated EP and OP cultures (Figure 5G). Separately, PRKCA loss increased CellROX signal, most prominently in EP and OP cultures, supporting a role for PKCα in oxidative-stress control during persistence (Figure 5H). To assess whether increased ROS, a consequence of PKCα downregulation, reduces persistence, N-acetyl-L-cysteine (NAC) was used to counteract PKCα loss or inhibition in osimertinib-treated HCC827 cells. The results demonstrated that NAC partially reversed the effects of *PRKCA* KO or enzastaurin co-therapy in TKI-treated cells, resulting in a roughly 2-fold increase in drug-tolerant clones, reaching levels comparable to single-agent osimertinib in HCC827 or HCC827 EV cells (Figures 5I and 5J). Collectively, these data indicate that PKCα-dependent *ALDH1A1* expression constrains ROS within persisters and thereby promotes drug tolerance, as antioxidant supplementation partially rescues clonogenic survival when PKCα is lost or inhibited.

### 2.7 PKCα enables a pre-existing CD44^High^ stem-like subpopulation to seed persistence

Mesenchymal states and enhanced cell motility are key functional manifestations of phenotypic plasticity and stemness.^36^ Our results implicating PKCα in the expression of *ALDH1A1*, a marker and functional regulator of CSCs, in TKI-induced cells, as well as reports suggesting PKCα to be a critical signaling modality in breast CSCs,^37^ led us to ask whether PKCα contributes selectively to TKI-induced phenotypic plasticity within a CSC-like subpopulation, rather than mediating tolerance uniformly across the entire persister pool. For this, we used the well-established CSC marker CD44 to sort putative stem-like populations in the erlotinib and osimertinib persister cultures.^38^ In line with previous reports, HCC827 EP and OP cells exhibited a CD44-high subpopulation that was absent in non-persister cells (Figures 6A and 6B). To study the properties of these distinct subpopulations, HCC827 EP and OP were sorted based on CD44 status, denoted CD44^−^ or CD44^+^ for low and high compartments, respectively (Figures 6C, 6D, and S5C). CD44^+^ cells displayed more efficient tumorsphere formation (Figure 6E), supporting their stem-like identity.^39,40^ In addition, cells derived from the CD44^+^ pool exhibited elevated *PRKCA* expression and activity, increased vimentin with loss of E-cadherin, and exclusive expression of *ALDH1A1* compared with CD44^−^ cells (Figures 6F, 6G, and 6H). Collectively, these results indicate that TKI-induced persisters include a CD44^+^ stem-like fraction defined by coordinated PKCα activation, EMT-associated features, and *ALDH1A1* expression.

**Figure 6.**
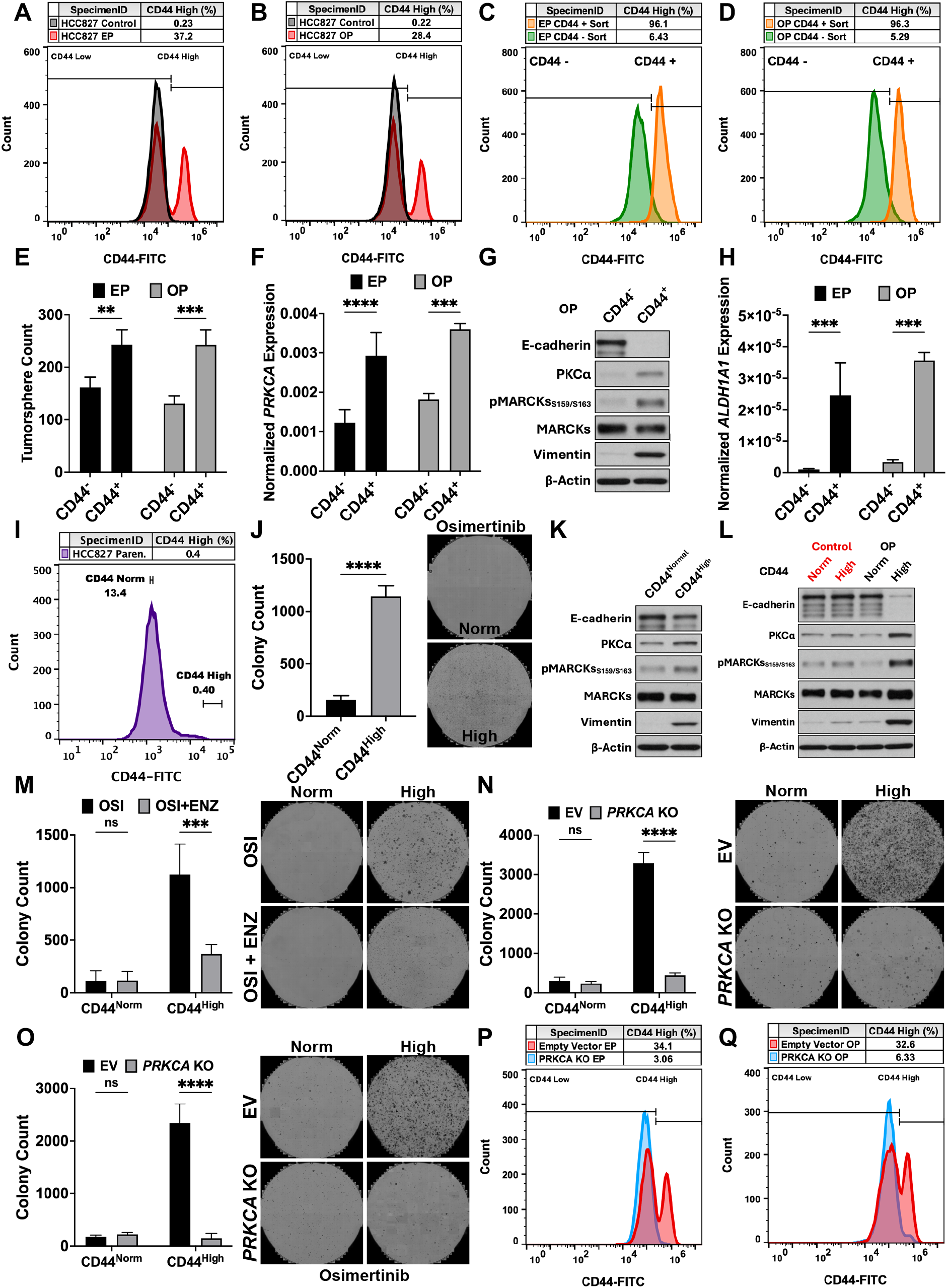
PKCα is necessary for survival of a pre-exisMng CD44^High^ compartment that seeds persistence. **(A**,**B)** Flow cytometry analysis of HCC827 control and EP **(A)** or OP **(B)** cells stained with CD44–FITC. **(C**,**D)** Flow cytometry analysis and validation of EP **(C)** or OP **(D)** cells sorted based on CD44 expression; the two sorted populations were labeled CD44^−^ and CD44^+^ (low vs high CD44 expression, respectively). **(E)** Tumorsphere formation of CD44^−^ and CD44^+^ sorted EP and OP cells (*n* = 3). **(F)** Mean normalized *PRKCA* expression (to *ACTB*) in CD44^−^ and CD44^+^ sorted EP and OP cells (*n* = 3). **(G)** Immunoblot depicting levels of PKCα, phospho-MARCKS_S159/S163_, and EMT markers in CD44^−^ and CD44^+^ sorted OP cells (*n* = 3). **(H)** Mean normalized *ALDH1A1* expression (to *ACTB*) in CD44^−^ and CD44^+^ sorted EP and OP cells (*n* = 3). **(I)** Flow cytometry analysis of treatment-naïve parental HCC827 cells. Brackets indicate sorted ranges labeled CD44^Norm^ (mean expression) and CD44^High^ (top 0.4% of CD44-expressing cells). **(J)** Clonogenic survival of treatment-naïve CD44^Norm^ and CD44^High^ cultures treated with OSI (10 nM) as described in 2A. Representative images of stained plates are shown (*n* = 3). **(K)** Immunoblot depicting levels of PKCα, MARCKS_S159/S163_, and EMT markers in CD44^Norm^ and CD44^High^ cultures (*n* = 3). **(L)** Immunoblot depicting levels of PKCα, MARCKS_S159/S163_, and EMT markers in CD44^Norm^ and CD44^High^ cultures treated with vehicle (DMSO) or OSI (10 nM) for 14 days, followed by a 10-day recovery (*n* = 2). **(M)** Clonogenic survival of CD44^Norm^ and CD44^High^ cells treated with OSI (10 nM) alone or OSI + ENZ (1 µM) for 14 days (*n* = 3). **(N**,**O)** Clonogenic survival of CD44^Norm^ and CD44^High^ cells expressing EV or with *PRKCA* KO treated with ERL (100 nM) **(N)** or OSI (10 nM) **(O)**. Representative images of stained plates are shown (*n* = 3). **(P**,**Q)** Flow cytometry analysis of HCC827 EP **(P)** or OP **(Q)** cells expressing EV or with *PRKCA* KO. Data are presented as mean ± SD. *p-va*lues for E,F,H, and J were determined by multiple unpaired two-sided t-tests. *p-va*lues for M–O were determined by two-way ANOVA with Šidák’s correction. \**p* < 0.05, \*\**p* < 0.01, \*\*\**p* < 0.001, \*\*\*\**p* < 0.0001, ^ns^*p* > 0.05.

With these findings, a key question emerged: whether the CD44^+^ population arises from a pre-existing CD44-high compartment in parental HCC827 cells. To address this, treatment-naïve HCC827 cells were sorted by CD44 expression to isolate two populations, one representative of the mean (CD44^Norm^), and one from the top 0.4% (CD44^High^) of CD44-expressing cells (Figure 6I). Interestingly, cultured CD44^High^ cells were intrinsically mesenchymal and more tolerant to osimertinib, exhibiting improved capacity for clonogenic survival relative to CD44^Norm^ (Figures 6J and S5D). However, in contrast to EP- and OP-derived persister fractions (CD44^−^ and CD44^+^), treatment-naïve CD44^High^ cells showed minimal differences in PKCα expression and activity compared with CD44^Norm^, and although vimentin was increased, E-cadherin was largely retained (Figure 6K). We therefore asked whether CD44^High^ cells exhibit enhanced phenotypic plasticity upon TKI exposure, explaining the observed variations between CD44^High^ and EP- or OP-derived CD44^+^. To this end, the sorted treatment-naïve CD44^Norm^ and CD44^High^ populations were treated with osimertinib or vehicle for 2 weeks to generate persister cultures. The results showed that CD44^High^ cells underwent substantially greater TKI-induced reprogramming, including increased PKCα and vimentin expression and near-complete loss of E-cadherin, changes that were not observed in CD44^Norm^ cells following osimertinib induction (Figure 6L). To test whether PKCα contributes to persister genesis selectively within the CD44^High^ pool, sorted CD44^Norm^ and CD44^High^ cells were treated with TKIs alone or with enzastaurin, or under conditions of PKCα loss. Here, CD44^Norm^ cells formed negligible colonies following TKI treatment, and suppressing PKCα in this pool did not measurably affect clonogenic survival (Figures 6M, 6N, 6O, and S5E). In contrast, CD44^High^ cells showed markedly increased generation of persistence, and, importantly, loss or inhibition of PKCα in these parental CD44^High^ cells markedly attenuated persistence, reducing the number of erlotinib- and osimertinib-tolerant clones (Figures 6M, 6N, and 6O). Consistent with this, *PRKCA* loss prevented enrichment of the CD44^High^ subpopulation during erlotinib- or osimertinib-induced persistence (Figures 6P and 6Q). Together, these findings support a model in which a rare, pre-existing CD44^High^ population serves as a principal substrate for persister formation and requires PKCα to execute the EMT and *ALDH1A1*-associated program, driving drug tolerance under EGFR TKI pressure.

## 3 Discussion

Drug-tolerant persister cells are proposed as a central barrier to durable therapeutic responses across cancers, forming reservoirs of residual cells that survive otherwise effective treatment.^11^ Biologically, persisters typically reflect early, reversible tolerance states, often involving dormancy and adaptive reprogramming rather than stable, mutation-defined resistance, and therefore represent a major therapeutic opportunity.^13^ These programs intersect strongly with phenotypic plasticity and CSC biology, including enrichment of stem-like subpopulations and mesenchymal features that support survival and outgrowth under drug pressure.^41–43^ A recurring feature of persistence is redox adaptation, in which persister cells engage antioxidant and metabolic mechanisms to buffer therapy-associated oxidative stress, limiting ROS and detoxifying lipid peroxidation products.^12,44,45^ To date, a major gap remains in defining the mechanisms that govern persister genesis and in translating these vulnerabilities into therapeutic strategies.^32^ Here, we position PKCα as a key effector required for the survival and outgrowth of a niche population of cells with strong attributes of EMT and stemness.

More specifically, our investigation identified PKCα as a critical modulator of early, reversible drug tolerance, as loss or inhibition of the kinase drastically suppressed the genesis of TKI-induced persister cells and significantly delayed relapse of tumors *in vivo*. Drug tolerance in our model was characterized by increased PKCα mRNA and protein, with mEGFR-independent activation of PKCα driving MARCKS and S6K phosphorylation. Given the negligible effects of PKCα inhibition on already established resistant cells, our data strongly indicate that PKCα plays a defining role in the early priming of treatment-naïve cells for survival and presents a unique therapeutic opportunity.

A central advance of this study is that it localizes the functional action of PKCα to a rare, pre-existing CD44^High^ stem-like subpopulation, identifying this minor persister-competent reservoir, instead of the bulk tumor population, as the critical substrate for TKI-tolerant survival and relapse. While PKCα was dispensable for basal growth in treatment-naïve cells, it was required for survival of residual cells after prolonged, but not acute, TKI exposure, implicating PKCα in drug tolerance and revealing a context-dependent role. However, rather than supporting a model in which PKCα broadly drives tolerance across the population, our findings indicate that persistence is largely biased toward a rare, pre-existing CD44^High^ compartment that is intrinsically more tolerant and uniquely competent for TKI-induced reprogramming. Specifically, treatment-naïve CD44^High^ cells exhibit greater clonogenic survival under osimertinib, and, upon TKI exposure, undergo markedly stronger transcriptional and phenotypic remodeling, inducing PKCα, increasing vimentin, and extinguishing E-cadherin, an effect that was not reproduced in CD44^Norm^ pools. Additionally, persisters demonstrated PKCα-dependent EMT reprogramming, as the kinase was required for induction, but not maintenance, of mesenchymal states in persister cells. Consistent with evidence that PKCα signaling promotes EMT-associated cellular plasticity and supports cancer stem-like phenotypes,^19,37,46^ persister cultures became enriched for a CD44^+^ tumorsphere-forming fraction with elevated PKCα activity, EMT features, and robust PKCα-dependent *ALDH1A1* expression. Importantly, PKCα suppression selectively compromises the persister-competent state, with loss or inhibition of the kinase markedly reducing clonogenic persistence arising from CD44^High^ cells and preventing enrichment of the CD44^High^ pool during TKI treatment, whereas CD44^Norm^ cells rarely persist and are largely unaffected by PKCα suppression. Mechanistically, PKCα is required for the robust induction of *ALDH1A1* during persister genesis *in vitro* and *in vivo*, and *ALDH1A1* depletion increases ROS in persister cultures under oxidative stress conditions. In line with reports on ROS buffering being functionally relevant to tolerance,^10,47^ antioxidant supplementation (NAC) partially rescued clonogenic survival when PKCα was genetically ablated or pharmacologically inhibited, supporting a model in which PKCα-driven *ALDH1A1* induction promotes an antioxidant state that enables survival and subsequent outgrowth of the persister-competent CD44^High^ state.

This work defines a specific context in which PKCα inhibition may have therapeutic value. Despite years of research, clinical PKCα inhibitors have failed as single agents or in combination with other treatment regimens.^26,48-50^ This may reflect, in part, limited biomarker-driven patient selection (e.g. as first happened with TKIs in NSCLC), contradictory roles of PKCα across cancer contexts, and the lack of isoform-specific inhibitors which further complicate the development of effective therapeutic strategies.^24,25^ The current setting provides a specific rationale for PKCα inhibition. Currently, persister generation is proposed as an important mechanism through which residual cells survive therapy and later acquire or select for secondary resistance mutations that enable clonal recovery.^7^ This remains a key problem in mEGFR-positive lung cancer where relapse after EGFR TKI therapy, including osimertinib-based regimens, remains a major challenge. Therefore, we propose that the addition of PKCα inhibitors to the current therapeutic regimen as an upfront combination therapy may be beneficial in delaying relapse as demonstrated in our *in vivo* preclinical models. Notably, our results indicate that the robust therapeutic effects of inhibiting PKCα are confined to the early phases of treatment, consistent with the timing-dependent effects observed *in vitro*.

Collectively, these results support a model in which PKCα functions as a context-dependent effector that enables survival and outgrowth of a pre-existing, plastic CD44^High^ subpopulation, potentially in part through *ALDH1A1*-mediated control of oxidative stress and thereby seeding drug-tolerant persistence under TKI pressure. Future studies should explore the molecular mechanisms underlying PKCα-driven persistence, particularly in relation to metabolic reprogramming and epigenetic modifications. Moreover, the apparent uncoupling of PKCα activation from mEGFR suggests a more general role in persistence, supported by preliminary unpublished observations in KRAS-mutant cells treated with sotorasib and pulmonary neuroendocrine cells treated with cisplatin. Hence, expanding the current model to assess PKCα-mediated drug tolerance in other treatment regimens is warranted.

## Acknowledgments

This work was supported by Merit Review Award I01 BX004621 from the United States Department of Veterans Affairs Biomedical Laboratory Research and Development Service. The authors wish to acknowledge the facilities at Stony Brook University, including the Division of Laboratory Animal Resources, the genomics core facility, the histology core, and flow cytometry core facility for their assistance and services. The schematics in this paper were generated using BioRender.

## Author Contributions

Conceptualization, M Sadeghi and Y.A.H.; Methodology, M Sadeghi, M Salama, and Y.A.H.; Software, M Sadeghi and J.Y.; Formal Analysis, M Sadeghi and J.Y.; Investigation, M Sadeghi, M Salama, A.H., and S.C.; Visualization, M Sadeghi; Writing – Original Draft, M Sadeghi; Writing – Review & Editing, M Salama, S.C., and Y.A.H.; Supervision, Y.A.H.; Funding Acquisition, Y.A.H.

## Declaration of Interests

The authors declare no competing interests.

## 4. Materials and Methods Reagents

Anti-ALDH1A1 (Cat#36671S and Cat#54135S [IHC]), p-AKT_T308_ (Cat#4056), p-AKT_S473_ (Cat#4060), E-cadherin (Cat#3195S), p-EGFR_Y1068_ (Cat#3777), p-EGFR_Y1173_ (Cat#4407S), EGFR (Cat#4267), p-ERK_T202/Y204_ (Cat#9101), ERK (Cat#4695), p-MARCKS_S152/156_ (Cat#2741), p-MARCK_S159/163_ (Cat#11992), N-cadherin (Cat#13116), p-P70S6K_T389_ (Cat#9234), P70S6K (Cat#2708), PKCα (Cat#2056), Vimentin (Cat#4650S and Cat#5741S [IHC]), and ZEB1 (Cat#70512S) antibodies were from Cell Signaling (Danvers, MA). Anti-AKT (Cat#44-609G), CD44-FITC (Cat#11-0441-86), Alexa Fluor™ 488 (Cat#A11008), p-MARCKS_S163_ (Cat#PA5-36852) and MARCKS (Cat#PA5-84812) antibodies were from Thermo Fisher (Waltham, MA). p-EGFR_Y992_ (Cat#AP0026) was from ABconal Technology (Woburn, MA). β-actin (Cat#A5441) and pEGFR_T654_ (Cat#04-282) were from Sigma Aldrich (St. Louis, MO). Rabbit IgG HRP (Cat#102645-182) and Mouse IgG HRP (Cat#102646-160) were from VWR International (Radnor, PA). 2× Laemmli Sample Buffer (Cat#1610737), Acacia (Cat#S25114), Formalin (Cat#SF1004), Nitro Blue Tetrazolium Chloride (Cat#N6495), Puromycin (Cat#NC9138068), Trypan Blue Solution (Cat#15250061), Bacteriological Agar (Cat#97064-336), Luria Broth Base (Cat#90003-252), PBS (Cat#97062-338), and Thiazolyl Blue Tetrazolium Bromide (Cat#97062-380) were from VWR International. RPMI-1640 (Cat#11875093), DMEM (Cat#11965092), D-PBS (Cat#14190250), Opti-MEM (Cat#31985062), FBS (Cat#26400044), PenStrep (Cat#15140122), RNA_iMAX_ (Cat#13778150), and Lipofectamine 2000 (Cat#11668019) were from Life Technologies. Enzastaurin (Cat#E-4506), Erlotinib (Cat#E-4997), and Osimertinib (Cat#O-7233) were from LC Laboratories (Woburn, MA). Matrigel (Cat#354230) was purchased from Corning (Corning, NY) and DAPI (Cat#F6057), crystal violet (Cat#C0775), and N-Acetyl-L-cysteine (Cat#A9165) were from Sigma-Aldrich. Human serum (Cat#009-000-121) was obtained from Jackson Laboratories (West Grove, PA). Critical commercial assays including PureLink RNA Kits (Cat#12183018A), SuperScript III (Cat#11752250), Fast-Advanced Master Mix (Cat#4444965), CellROX Green (Cat#C10444), TaqMan Assays, and siRNA were purchased from Thermo Fisher or Qiagen (refer to Table S2).

### Cell lines

HCC827, HCC4006, and HEK293T were purchased from American Type Culture Collection (Manassas, VA). H1975 and PC9 cells were kind gifts from Jason Sheltzer PhD at Cold Spring Harbor. H3255 cells were obtained from Dr. Joseph Contessa at Yale School of Medicine. HCC827, HCC4006, H3255, PC9, and H1975 were grown in RPMI supplemented with 10% (v/v) fetal bovine serum (FBS) and 1% PenStrep. HEK293T was maintained in DMEM with 10% (v/v) FBS and 1% PenStrep. Cell lines were cultured in a humidified incubator at 37°C with 5% CO_2_ and regularly tested for mycoplasma contamination. HCC827, the primary cell-line model used in this work, was authenticated by STR profiling. Resistant cells were generated through treatment with 100 nM erlotinib or 10 nM osimertinib over the course of 2 months in serum-deprived medium. siRNA transfections were performed using Lipofectamine RNAi_MAX_ and 20 nM siRNA according to the manufacturer’s protocol. Transfected cells were grown in medium with 10% FBS for 48 hours then seeded for further experiments as indicated.

### Animal models

All *in vivo* studies were done in compliance with protocols approved by Institutional Animal Care and Use Committee (IACUC) at Stony Brook University. Six-week-old female NU/J (Foxn1^nu/nu^) or 8-week-old female NSG (Prkdc^scid^ Il2rg^tm1Wjl^/SzJ) mice were obtained from Jackson Laboratories (Bar Harbor, ME) and allowed 10 days to acclimate prior to randomization and initiation of experiments. NSG mice arrived with subcutaneous TM00193 (EGFR_del746-750,_ treatment-naïve, obtained during surgical resection) tumors. Animals were housed in groups of 3–4 in pathogen-free conditions under constant temperature and humidity with 12-hour light and 12-hour dark cycles from 6 AM to 6 PM.

### CRISPR/Cas9-mediated generation of PKCα knockout cells

PKCα knockout (KO) HCC827 cells were generated using the lentiviral CRISPR/Cas9 v2 system: pLenti_CRISPR/Cas9v2_ (Addgene #52961). A guide targeting *PRKCA* (sg*PRKCA*) with the target sequence 5′-GAGGCAGAAGAACGTGCACG-3′ was cloned using the forward and reverse oligonucleotides: 5′-CACCGAGGCAGAAGAACGTGCACG-3′ and 5′-AAACCGTGCACGTTCTTCTGCCTC-3′, respectively. Lentiviral particles were produced by seeding 5.5 × 10^5^ HEK293T cells/well in a 6-well plate in DMEM + 10% FBS and transfecting the next day with 6.3 µL Lipofectamine 2000 diluted in 30 µL Opti-MEM together with plasmids diluted in 30 µL Opti-MEM: 1 µg pLenti_CRISPR/Cas9 v2–sgPRKCA_, 1 µg psPAX2 (Addgene #12260), and 100 ng pCMV-VSV-G (Addgene #8454). After overnight incubation, medium was replaced with DMEM + 30% FBS, and viral supernatant was collected 48 h post-transfection, filtered through a 0.45-µm PVDF filter, and stored at −80°C. HCC827 cells were plated at 2.0 × 10^5^ cells/well in 6-well plates in RPMI + 10% FBS and infected the following day in 1 mL RPMI + 10% FBS containing 16 µg/mL polybrene (28728-55-4; Santa-Cruz Biotechnology) by adding 1 mL lentiviral supernatant per well. Medium was replaced after 24 h, and cells were expanded 2 days post-infection into 100-mm dishes and selected with 5 µg/mL puromycin for 10 days prior to experimentation; empty vector control cells were generated in parallel.

### MTT, growth, and cell viability assays

To assay viable cell numbers, 1×10^5^ or 5×10^4^ cells were seeded in 6- or 12-well plates, respectively, and allowed to adhere overnight; the cells were then serum-starved for 5 hours and treated as indicated for 48 hours. Thiazolyl Blue Tetrazolium Bromide (MTT; 5 mg/mL in DPBS) and fresh medium were added in 1:1 mixture to each well and incubated at 37°C for 60 minutes. Media was aspirated and 1 or 2 mL (for 6- or 12-well plates respectively) DMSO was added to wells. Following brief agitation, optical density was read at 570 nm using SpectraMax M5 (Molecular Devices, Sunnyvale, CA). For cell viability assays, 1×10^5^ cells were seeded in 6-well plates overnight. Cells were then serum-starved and a 1:1 mixture of homogenized cell suspension from appropriate wells and trypan blue solution was prepared at the indicated time points. Live cells were counted using a hemocytometer and the total number of live cells at each indicated time point was recorded.

### Real-time qPCR

Total mRNA was extracted from 1×10^6^ cells using the PureLink RNA Mini Kit and converted to cDNA using the Superscript III Kit. Real-time qPCR was performed on the ABI 7500 Fast real-time system using the fast-advanced master mix and TaqMan assays provided by Thermo Fisher. The obtained C_t_ values were converted to mean normalized expression using the ΔΔ^Ct^ method with *ACTB* (β-actin) as the reference gene.

### Western blotting analysis

Gel electrophoresis was performed using Novex 4-20% gradient Midi-Gels run in Criterion™ Cell Tanks (#1656001, Bio-Rad) then transferred onto 0.45 µm nitrocellulose membranes using Criterion™ Blotter (#1704070, Bio-Rad). Membranes were incubated with primary antibodies overnight at 4°C. Blots were incubated with HRP-conjugated secondary antibodies and detected with Pierce™ ECL reagent.

### Transwell migration assay

For migration, 2.5×10^4^ cells were seeded in transwell inserts (8 µm pores) in serum-free medium. Cells were allowed 16 hours to migrate following the addition of FBS-supplemented medium (10%) to the outer well. The inserts were then fixed with 70% ethanol, stained with 0.5% crystal violet, and swabbed to remove non-migrated cells from the upper surface of the membrane. To quantitate migrating cells, inserts were imaged and quantified in ImageJ.

### Clonogenic survival assay and generation of persisters

To induce drug-tolerant clones, 2×10^6^ cells were seeded in 100 mm dishes and treated with the appropriate inhibitor or combination of inhibitors for 14 days in serum-deprived medium. Surviving clones were then allowed 7 to 14 days for regrowth in drug-free and serum-supplemented medium to form colonies. The colony counts per treatment condition were quantitated as a measurement of the number of original cells that survived within each treatment condition. For quantitation, cells were fixed with 100% methanol for 10 minutes, stained using 0.5% crystal violet for 20 minutes, washed, and imaged using the Nexcelom Celigo Image Cytometer. Monoclonal persisters were generated by seeding 6×10^6^ cells in 150 mm dishes. The cells were treated the next day with vehicle or the appropriate inhibitor in serum-free medium for 14 days. Surviving cells were replated in a 100 mm dish with moderate agitation to ensure a single cell suspension. Cells were then allowed to grow for roughly 7 days in drug-free and serum-supplemented medium. Isolated colonies of persisters were then transferred to a 24-well plate for propagation. In the case of generating polyclonal persisters, all colonies were harvested and passaged in a new 100 mm dish. To generate the appropriate controls for persister experiments, seeded cells were treated with DMSO alongside experimental conditions for 14 days in serum-deprived medium, followed by a 7-day regrowth in serum-supplemented medium.

### Immunofluorescence

3×10^5^ cells were seeded in 35-mm glass-bottom dishes (poly-D-lysine coated). The next day, cells were serum-starved for 16 hours overnight then treated for 30 minutes with the appropriate TKI. Cells were fixed using 3.7% paraformaldehyde followed by permeabilization with 100% ice-cold methanol, both with 10-minute incubation times. Following blocking for 1 hour in 2% human serum, cells were incubated with the appropriate primary antibody overnight at 4°C. Samples were incubated with fluorophore-conjugated secondary antibodies at room temperature for 1 hour prior to the addition of mounting medium containing DAPI. Samples were imaged using a Leica TCS SP8 confocal microscope system.

### Reactive Oxygen Species (ROS) Detection

Intracellular reactive oxygen species (ROS) were assessed using CellROX Green Reagent (#C10444, Thermo Fisher Scientific) and quantified by whole-well fluorescence imaging of live cells. Erlotinib persister (EP) and osimertinib persister (OP) cultures were generated as described above and included EP/OP derived from empty vector (EV) control or *PRKCA* knockout (KO) cells, as well as HCC827 EP and OP cultures transfected with si*ALDH1A1*. Cells were plated at 5.0 × 10^4^ cells/well in 12-well tissue culture plates and allowed to adhere. Prior to ROS induction, cells were serum-starved for 5 h, then treated with 200 µM menadione (Cat# HY-B0332; MedChemExpress) for 1 h at 37°C. Following menadione exposure, cells were incubated with CellROX Green at a final concentration of 5 µM for 30 min at 37°C, protected from light. Cells were then washed three times with D-PBS and immediately imaged using a Celigo imaging cytometer (Nexcelom) using the green fluorescence channel with identical acquisition settings across conditions. ROS levels were quantified from CellROX Green fluorescence intensity (background-subtracted) and reported relative to the appropriate matched controls.

### Flow cytometry

A suspension of 1×10^7^ cells was prepared in FACS buffer (1× PBS, 2% FBS, 2 mM EDTA, and 0.1% sodium azide). The cell suspension (100 µL) was transferred to round bottom tubes and incubated with the appropriate fluorophore-conjugated primary antibody at room temperature for 30 minutes. Cells were then washed twice and resuspended in 200–300 µL FACS buffer. Samples were kept on ice until analysis using BD Accuri™ C6 Plus Flow Cytometer. For cell sorting based on CD44 expression levels, 5×10^7^ cells were suspended in D-PBS supplemented with 10% FBS. Following incubation with FITC-conjugated primary antibody, cells were washed 3 times before resuspension in FBS supplemented D-PBS and sorted using FACSAria IIIu Cell Sorter (BD Biosciences).

### Tumorsphere formation assay

To assay tumorsphere formation, 0.4% agar was prepared by mixing 2× RPMI medium supplemented with 2% (v/v) FBS 1:1 with sterile 0.8% agar. A small volume (>200 µL) of medium containing 1×10^5^ cells was added to the warm 0.4% agar, which was then plated in ultra-low attachment 6-well plates and allowed to cool for 45–60 min to solidify. Plates were incubated for 2–4 weeks at 37°C. Fresh medium (200 µL; 1% FBS) was added to wells every 5 days. Cells were stained overnight with 200 µL Nitro Blue Tetrazolium Chloride (NBT; 1 mg/mL) upon the conclusion of the experiment. Plates were imaged and tumorspheres were counted using Nexcelom Celigo Image Cytometer.

### Xenograft studies

For all experiments, mice were allowed a 10-day acclimation period. For xenografting studies, immunodeficient (NU/J) mice were subcutaneously injected with a 1:1 mixture of 50 µL matrigel and 50 µL DPBS containing 5×10^6^ HCC827 parental, HCC827 EV, or *PRKCA* KO cells. In the case of patient-derived xenografts (PDXs), NSG mice were received with prior injections of TM00193 tumors. After growth of tumors to a volume of ~150 mm^3^, mice were treated with vehicle (10% acacia), erlotinib (40 mg/kg/day), osimertinib (25 mg/kg/day), enzastaurin (125 mg/kg twice daily), or an appropriate combination of inhibitors as indicated in the figure captions. For discontinuous studies, treatments were withheld at the indicated time points; otherwise, the inhibitors were administered until tumor relapse was observed in all groups (roughly 90 days). Inhibitors were suspended in 10% acacia and administered via oral gavage. Tumor measurements were taken every other day or weekly using a caliper. Tumor volumes were calculated using the formula: Volume = Length × 0.5 Width^2^. Tumors were excised at endpoint and fixed in 10% formalin. Formalin-fixed tissues were submitted to Research Histology Core Laboratory at Stony Brook University for paraffin embedding, sectioning, and H&E staining. Immunohistochemistry was performed based on standard protocols, with antigen retrieval and hematoxylin counterstaining. Slides were imaged at 5×–10× magnification using AXIO Imager M2 (Zeiss) equipped with Axiocam 208 Color.

### Quantification and statistical analysis

GraphPad Prism was used for data graphing and statistical calculations. FlowJo was used for flow cytometry analysis and visualizations. Data are presented as mean ± standard deviation (SD) unless otherwise noted, with biological replicate count (*n*) denoted in the figure legends. For statistical analysis, a *p* < 0.05 was considered significant. All statistical tests involved at least 3 biological replicates. An unpaired t test was performed for comparisons of two groups. Multiple t tests were followed by Holm-Šidák’s correction. For comparison of more than two groups, a one-way ANOVA was performed with Šidák’s correction. For comparisons involving two categorical variables a two-way ANOVA was used followed by Šidák’s multiple comparisons test. For longitudinal datasets with repeated measurements and missing values, data were analyzed in GraphPad Prism using a mixed-effects model with time, treatment condition, and the time × treatment interaction as fixed effects and subject as the repeated-measures factor; Geisser-Greenhouse correction was applied, and treatment conditions were compared at each time point using Šidák’s multiple-comparisons test with multiplicity-adjusted *p*-values reported.

## Notes

### Competing Interest Statement

The authors have declared no competing interest.

