## Supplemental Figures for "TKI-Tolerant Persisters Emerge from a PKCα-Dependent and Highly Plastic Subpopulation of Stem-Like Cells in NSCLC"


**Author Contributions:** Conceptualization, M Sadeghi and Y.A.H.; Methodology, M Sadeghi, M Salama, and Y.A.H.; Software, M Sadeghi and J.Y.; Formal Analysis, M Sadeghi and J.Y.; Investigation, M Sadeghi, M Salama, A.H., and S.C.; Visualization, M Sadeghi; Writing – Original Draft, M Sadeghi; Writing – Review & Editing, M Salama, S.C., and Y.A.H.; Supervision, Y.A.H.; Funding Acquisition, Y.A.H.

**Competing Interest Statement:** The authors declare no competing interests.


**Keywords:** Non-Small Cell Lung Cancer (NSCLC); Tyrosine Kinase Inhibitors (TKI); Drug-Tolerant Persisters (DTPs); Drug Resistance; Cancer Stem Cells; Epidermal Growth Factor Receptor (EGFR); Epithelial-to-Mesenchymal Transition (EMT); Protein Kinase C (PKC); Reactive Oxygen Species; ALDH1A1.


**This PDF file includes:**


Supplemental Figures S1 to S5
Tables S1 to S2

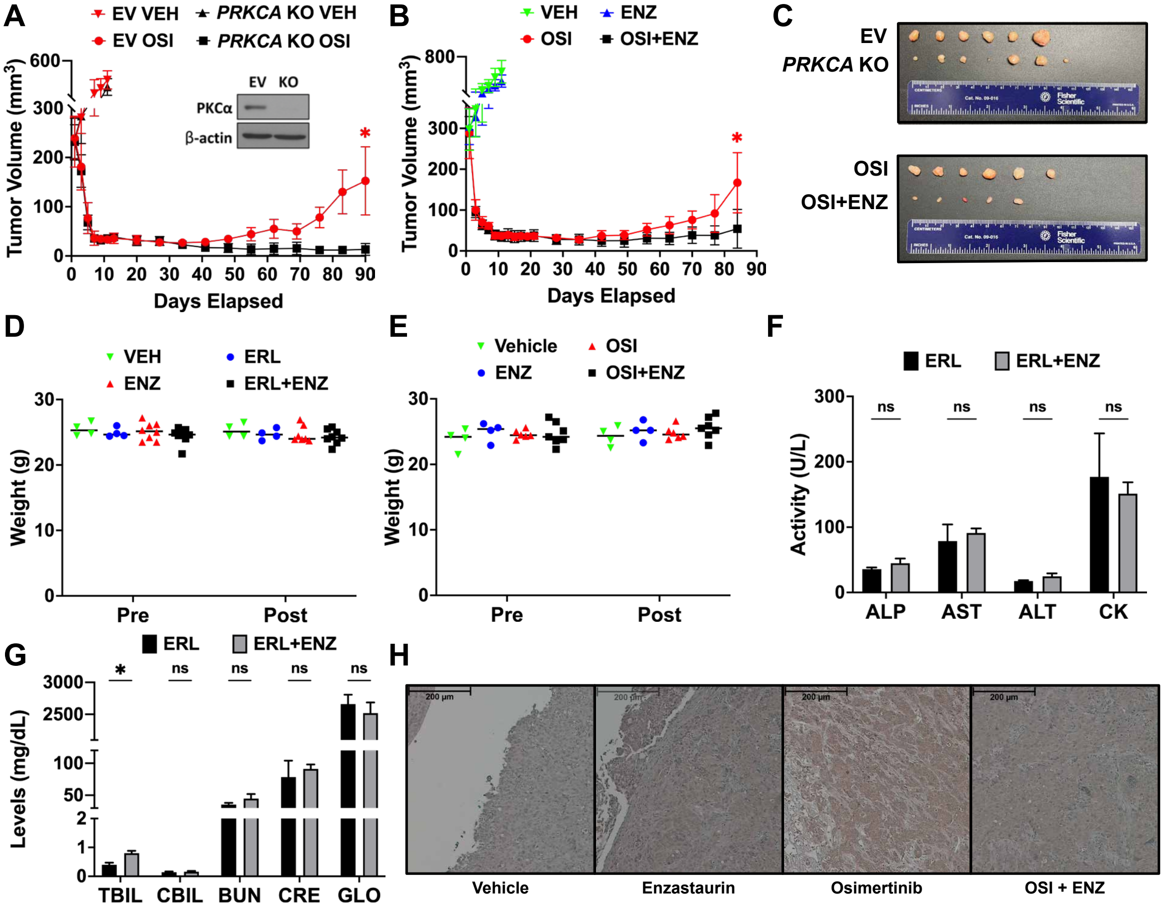


**Figure S1. Effects of inhibition or loss of *PRKCA* on relapse following TKI treatment**

(**A**) Tumor volumes of HCC827 empty vector (EV) and *PRKCA* knockout (KO) xenografts treated with vehicle (VEH; 10% acacia) or osimertinib (OSI; 25 mg/kg/day) over 90 days ($n$ = 6–7). Loss of PKCα in HCC827 validated by immunoblot as shown. (**B**) Tumor volumes of HCC827 xenografts treated with VEH (10% acacia), OSI (25 mg/kg/day), enzastaurin (ENZ; 125 mg/kg twice daily), or OSI in combination with ENZ over 83 days ($n$ = 6). (**C**) Images of excised tumors from panels A and B. (**D**) Mouse body weight measurements from Figure 1B taken before treatment initiation and after treatment conclusion (day 85). The weights of mice treated with VEH (10% acacia), erlotinib (ERL; 40 mg/kg/day), ENZ (125 mg/kg, twice daily), or ERL in combination with ENZ are reported ($n$ = 4–7). (**E**) Mouse body weight measurements from the cohort in panel B taken before treatment initiation and after treatment conclusion (day 83). The weights of mice treated with VEH, OSI (25 mg/kg/day), ENZ (125 mg/kg, twice daily), or OSI in combination with ENZ are reported ($n$ = 4–7). (**F**) Markers of liver function: alkaline phosphatase (ALP), aspartate aminotransferase (AST), and alanine aminotransferase (ALT) and muscle health (creatine kinase; CK), and (**G**) levels of other appropriate indicators (Total bilirubin; TBIL, conjugated bilirubin; CBIL, blood urea nitrogen; BUN, creatinine; CRE, and globulin; GLO) for ERL (40 mg/kg/day) or ERL in combination with ENZ (125 mg/kg, twice daily) treated mice at endpoint from Figure 1B ($n$ = 5). (**H**) PKCα-immunostained sections obtained at endpoint (day 83) from HCC827 xenografts shown in panel B. Data are presented as mean ± SD. $p$-values in panels A and B were determined using a mixed-effects model. $p$-values in panels F and G were calculated using multiple unpaired, two-sided t tests. *$p < 0.05$, $^{ns}p > 0.05$ or not significant.

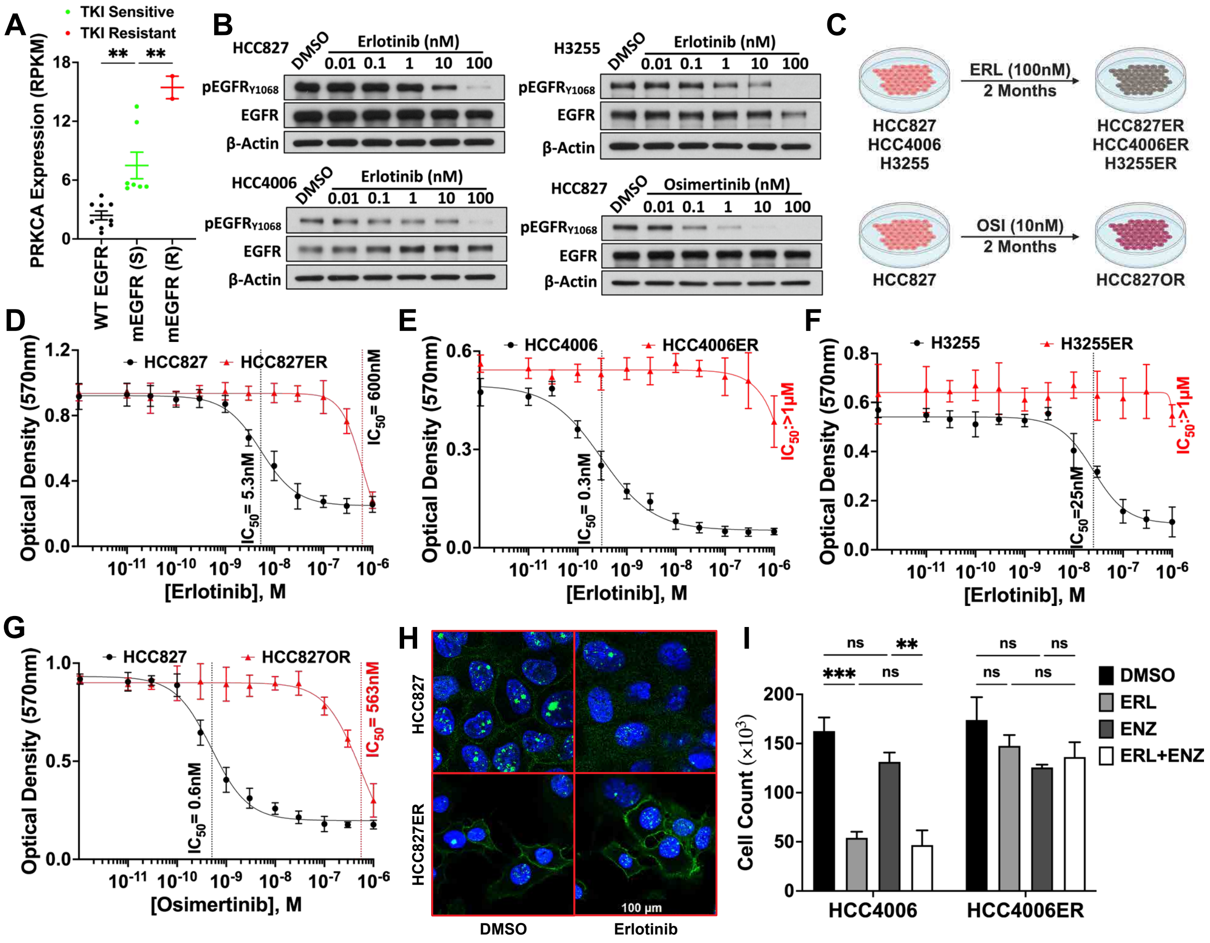


**Figure S2. Expression, activity, and the role of PKCα in TKI-insensitive cells**

**(A)** Expression of PKCα in cell lines harboring wild-type (WT) epidermal growth factor receptor (EGFR) or mutant EGFR (mEGFR; $EGFR_{del746-750}$ or $EGFR_{L858R}$) that are sensitive (S) or resistant (R) to TKI. Data were retrieved from the Cancer Cell Line Encyclopedia[1] ($n$ = 2–9). **(B)** Immunoblot assessing phosphorylation of $EGFR_{Y1068}$ in HCC827, HCC4006, or H3255 parental cells in response to the indicated doses of erlotinib (ERL) or osimertinib (OSI) following a treatment duration of 6 h ($n$ = 2). **(C)** Schematic depicting the doses and duration of treatment for the generation of resistant cell lines used in this study. **(D)** ERL dose response of HCC827 parental and HCC827ER (erlotinib-resistant) cells or **(E)** HCC4006 parental and HCC4006ER cells or **(F)** H3255 parental and H3255ER cells were treated with varying doses of ERL for 48 h. Optical density measurements were then made following 60 min incubation with MTT (thiazolyl blue tetrazolium bromide) ($n$ = 3). **(G)** Dose response to OSI of HCC827 parental and HCC827OR (osimertinib-resistant) cells. Cells were treated with varying doses of OSI for 48 h, then quantified as described in panels D–F ($n$ = 3). **(H)** Assessment of membrane localization of PKCα (green) in HCC827 parental and HCC827ER cells treated with dimethyl sulfoxide (DMSO) or ERL (100 nM) ($n$ = 2). **(I)** Cell counts of HCC4006 and HCC4006ER cells treated with DMSO, ERL (100 nM), enzastaurin (ENZ; 1 μM), or ERL in combination with ENZ ($n$ = 3). Data are presented as mean ± SD. $p$-values in panel A were determined by one-way ANOVA with Šidák's multiple comparisons test. The $p$-values in panel I were determined by two-way ANOVA with Šidák's multiple comparisons test. *$p < 0.05$, **$p < 0.01$, ****$p < 0.0001$, $^{ns}p > 0.05$ or not significant.

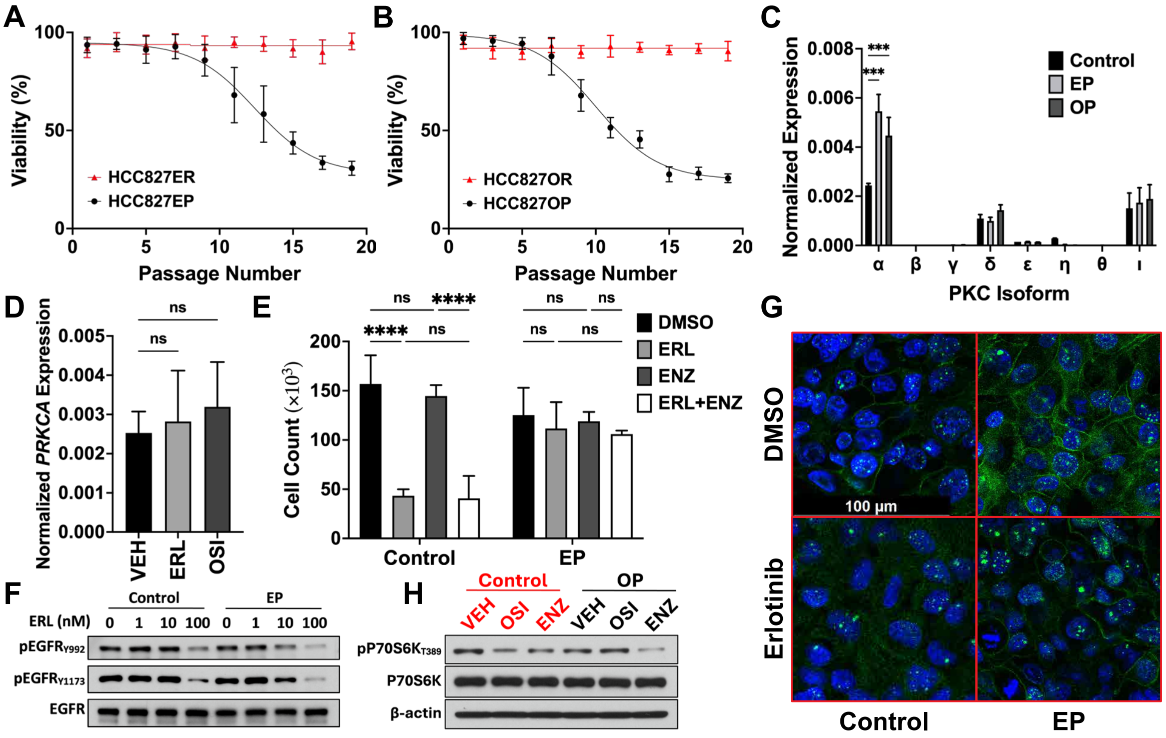


**Figure S3. Effects of passage number and PKC inhibition on the resensitization of persisters**

**(A)** Sensitivity of HCC827 erlotinib persisters (EP) and HCC827ER (erlotinib-resistant) cells to erlotinib (ERL) or **(B)** HCC827 osimertinib persisters (OP) and HCC827OR (osimertinib-resistant) cells to osimertinib (OSI) over 19 passages. Sensitivity was assessed every 2 passages by treating cells with dimethyl sulfoxide (DMSO) or the relevant tyrosine kinase inhibitor (TKI) for 48 h (EP/HCC827ER: ERL, 100 nM; OP/HCC827OR: OSI, 10 nM). Following MTT (thiazolyl blue tetrazolium bromide) treatment, viability was quantified by reporting the optical density of TKI-treated samples as a fraction of the DMSO control ($n$ = 3). **(C)** Normalized expression (to *ACTB*) of different PKC isoforms in control, EP, and OP cells ($n$ = 3). **(D)** Expression of *PRKCA* (encoding PKCα) in HCC827 parental cells following 24 h treatment with DMSO, ERL (100 nM), or OSI (10 nM) ($n$ = 3). **(E)** Cell counts of HCC827 control and EP cells treated with DMSO, ERL (100 nM), enzastaurin (ENZ; 1 µM), or ERL in combination with ENZ ($n$ = 3). **(F)** Immunoblot showing phosphorylation of EGFR at two sites that couple to PKC signaling in control and persister cells following ERL treatment at varying doses ($n$ = 2). **(G)** PKCα membrane localization (green) in HCC827 control and EP cells following ERL treatment (100 nM) ($n$ = 3). **(H)** Immunoblot depicting phosphorylation of $p70S6K_{T389}$ in control or OP cells, with treatment of DMSO (VEH), 10 nM OSI, or 1 µM ENZ ($n$ = 2). Data are presented as mean ± SD. $p$-values in panel D were determined by one-way ANOVA with Šidák's multiple comparisons test. $p$-values in panels C and E were determined by two-way ANOVA with Šidák's multiple comparisons test. **$p < 0.01$, ***$p < 0.001$, ****$p < 0.0001$, $^{ns}p > 0.05$ or not significant.

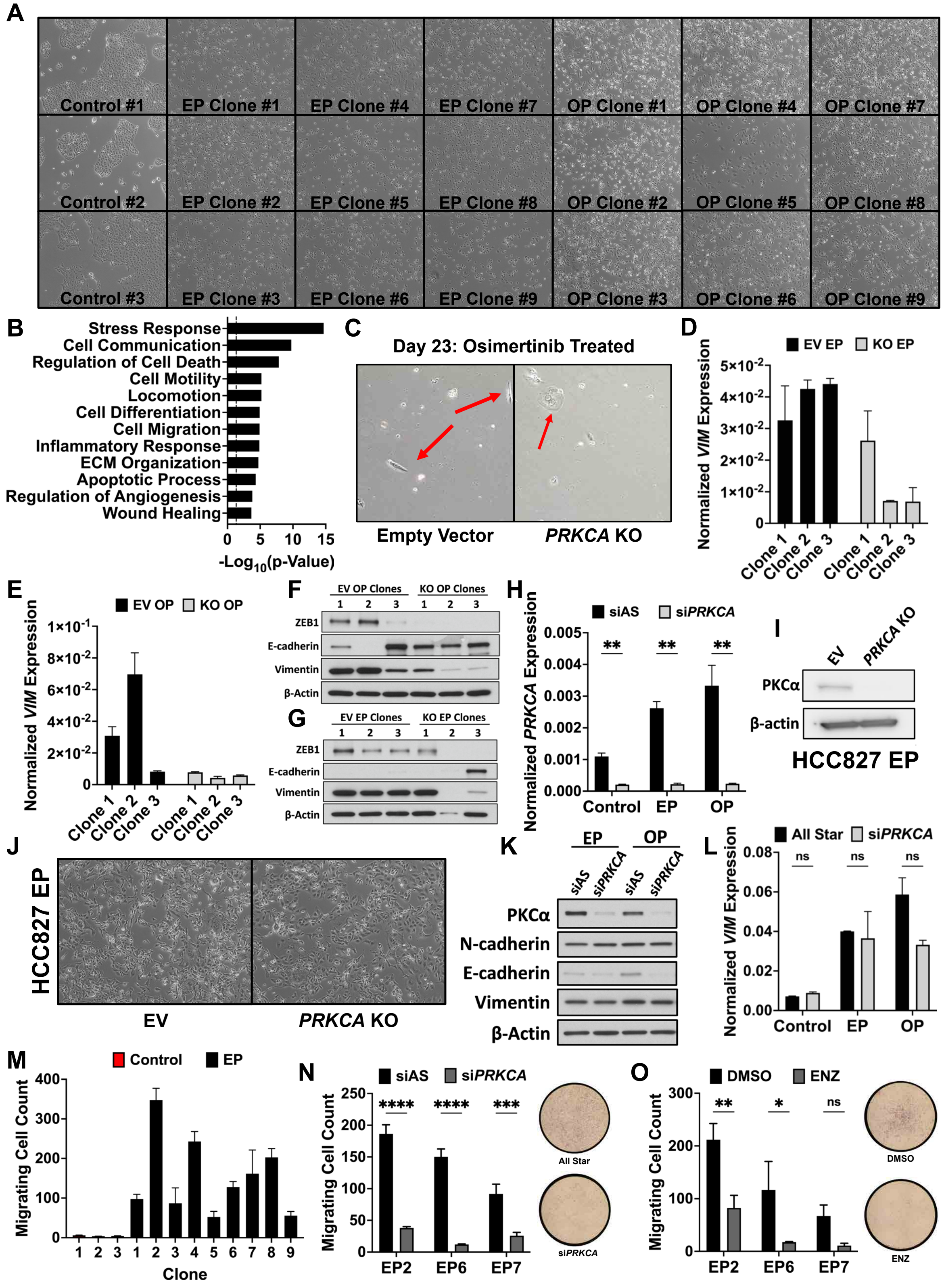


**Figure S4. Effect of PKCα suppression on EMT and migration of persisters**

**(A)** Bright-field microscopy images of HCC827 control, erlotinib persister (EP) clones (EP1–EP9), or osimertinib persister (OP) clones (OP1–OP9). Images were taken under 10× magnification. **(B)** Gene Ontology analysis of differentially expressed genes in HCC827 OP compared to control ($n$ = 3). **(C)** Images of residual cells at the 23-day time point from Figure 1G. Residual cells are generated following osimertinib treatment of empty vector (EV) and *PRKCA* knockout (KO) HCC827 cells. **(D-E)** Normalized expression (to *ACTB*) of *VIM* (Vimentin) in control and monoclonal erlotinib (EP) **(D)** or osimertinib persisters (OP) **(E)** generated from HCC827 empty vector (EV) and *PRKCA* knockout (KO) cells. **(F-G)** Western blot analysis of ZEB1, E-cadherin, and vimentin in EV and *PRKCA* KO monoclonal EP **(F)** or OP **(G)** ($n$ = 3). **(H)** Normalized expression of *PRKCA* in HCC827 control, EP, and OP cells following treatment with 20 nM All-Star control small interfering RNA (siAS) or 20 nM *PRKCA* small interfering RNA (si*PRKCA*) for 48 h ($n$ = 3). **(I)** Immunoblot validation of PKCα levels following CRISPR-mediated loss of *PRKCA* in HCC827 EP cells ($n$ = 2). **(J)** Bright-field microscopy images of HCC827 EP cells expressing an EV or with loss of *PRKCA* ($n$ = 2). **(K)** Immunoblot assessing the effects of *PRKCA* knockdown (KD) on the expression of vimentin, N-cadherin, and E-cadherin in EP and OP cells ($n$ = 2). **(L)** Normalized expression of *VIM* (vimentin) in HCC827 control, EP, and OP cells treated with siAS or *PRKCA* siRNA ($n$ = 3). **(M)** Transwell migration of HCC827 control and EP clones. Cells were allowed 16 h for migration to the bottom chamber, then quantified after staining and imaging via ImageJ ($n$ = 3). **(N)** Migrating cell counts of HCC827 EP clones pre-treated with 20 nM siAS or *PRKCA* siRNA ($n$ = 3). **(O)** Migrating cell counts of HCC827 EP clones pre-treated with dimethyl sulfoxide (DMSO) or enzastaurin (ENZ; 1 µM) for 6 h. Cells were then allowed 16 h to migrate in the presence of DMSO or ENZ ($n$ = 3). Data are presented as mean ± SD. $p$-values in panels H, N, and O were determined by multiple unpaired, two-sided t tests. $^{*}p < 0.05$, $^{**}p < 0.01$, $^{***}p < 0.001$, $^{****}p < 0.0001$, $^{ns}p > 0.05$ or not significant.

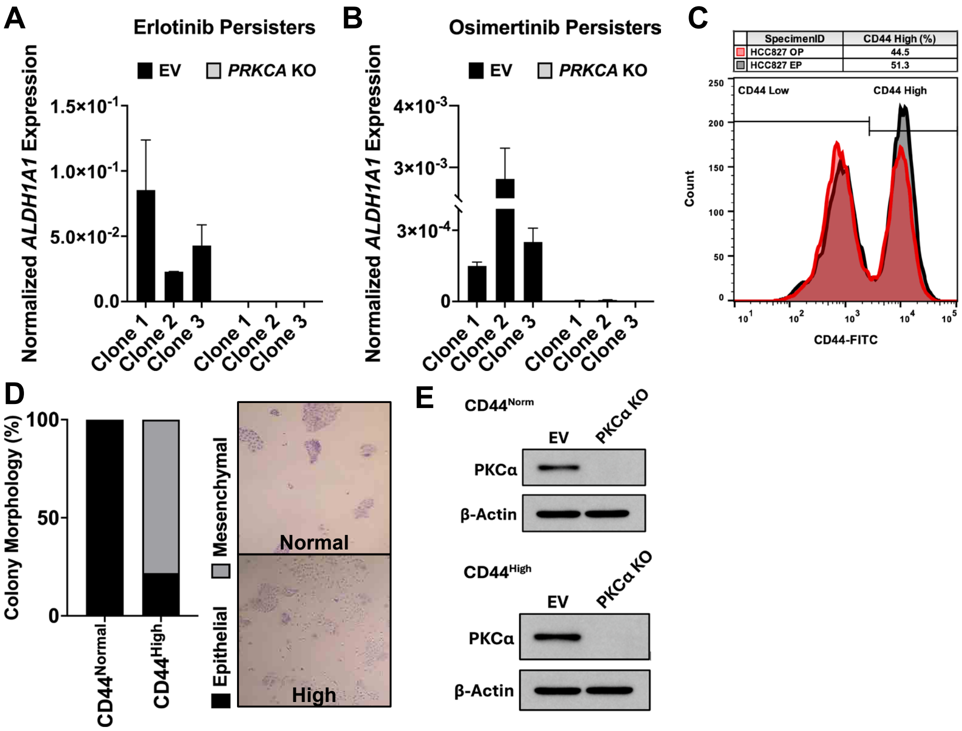


**Figure S5. Effects of *PRKCA* loss on *ALDH1A1* expression in persisters**

(**A)** Normalized expression of *ALDH1A1* in erlotinib or **(B)** osimertinib monoclonal persisters with preemptive loss of *PRKCA* (*n* = 3). **(C)** Flow cytometry analysis and validation of EP and OP cells. Brackets indicate sorted ranges labeled $CD44^-$ (low expression; bottom 50%) and $CD44^+$ (high expression; top 50%). **(D)** Quantification of the number of epithelial or mesenchymal colonies observed in treatment naïve $CD44^{Norm}$ and $CD44^{High}$ HCC827 cells, sorted as described in Figure 6I. States were assigned by measuring intercellular distances within colonies using ImageJ; predominantly non-zero distances were scored as mesenchymal. Representative images are shown (*n* = 10). **(E)** Immunoblot validating the loss of *PRKCA* in $CD44^{Norm}$ and $CD44^{High}$ HCC827 cells. Data are presented as mean ± SD.

**Table S1** EGFR Secondary Mutation Status in Xenograft Tumor Samples Following TKI Treatment

| | | | T790 | | | | | | | C797 | | | |
|---|---|---|---|---|---|---|---|---|---|---|---|---|---|
| **WT EGFR Sequence** | **C T C** | **A T C** | **A C G** | **C A G** | **C T C** | **A T G** | **C C C** | **T T C** | **G G C** | **T G C** | **C T C** | **C T G** | **G A C** |
| **Translation** | **L** | **I** | **T** | **Q** | **L** | **M** | **P** | **F** | **G** | **C** | **L** | **L** | **D** |
| **Protein Residue Number** | **788** | **789** | **790** | **791** | **792** | **793** | **794** | **795** | **796** | **797** | **798** | **799** | **800** |
| **Empty Vector Control #1** | C T C | A T C | A C G | C A G | C T C | A T G | C C C | T T C | G G C | T G C | C T C | C T G | G A C |
| **PRKCA KO Control #1** | C T C | A T C | A C G | C A G | C T C | A T G | C C C | T T C | G G C | T G C | C T C | C T G | G A C |
| **Empty Vector Erlotinib #1** | C T C | A T C | A C G | C A G | C T C | A T G | C C C | T T C | G G C | T G C | C T C | C T G | G A C |
| **Empty Vector Erlotinib #2** | C T C | A T C | A C G | C A G | C T C | A T G | C C C | T T C | G G C | T G C | C T C | C T G | G A C |
| **Empty Vector Erlotinib #3** | C T C | A T C | A C G | C A G | C T C | A T G | C C C | T T C | G G C | T G C | C T C | C T G | G A C |
| **Empty Vector Erlotinib #4** | C T C | A T C | A C G | C A G | C T C | A T G | C C C | T T C | G G C | T G C | C T C | C T G | G A C |
| **Empty Vector Erlotinib #5** | C T C | A T C | A C G | C A G | C T C | A T G | C C C | T T C | G G C | T G C | C T C | C T G | G A C |
| **PRKCA KO Erlotinib #1** | C T C | A T C | A C G | C A G | C T C | A T G | C C C | T T C | G G C | T G C | C T C | C T G | G A C |
| **PRKCA KO Erlotinib #3** | C T C | A T C | A C G | C A G | C T C | A T G | C C C | T T C | G G C | T G C | C T C | C T G | G A C |
| **PRKCA KO Erlotinib #4** | C T C | A T C | A C G | C A G | C T C | A T G | C C C | T T C | G G C | T G C | C T C | C T G | G A C |
| **PRKCA KO Erlotinib #5** | C T C | A T C | A C G | C A G | C T C | A T G | C C C | T T C | G G C | T G C | C T C | C T G | G A C |
| **PRKCA KO Erlotinib #8** | C T C | A T C | A C G | C A G | C T C | A T G | C C C | T T C | G G C | T G C | C T C | C T G | G A C |
| **Empty Vector Osimertinib #1** | C T C | A T C | A C G | C A G | C T C | A T G | C C C | T T C | G G C | T G C | C T C | C T G | G A C |
| **Empty Vector Osimertinib #2** | C T C | A T C | A C G | C A G | C T C | A T G | C C C | T T C | G G C | T G C | C T C | C T G | G A C |
| **Empty Vector Osimertinib #3** | C T C | A T C | A C G | C A G | C T C | A T G | C C C | T T C | G G C | T G C | C T C | C T G | G A C |
| **Empty Vector Osimertinib #4** | C T C | A T C | A C G | C A G | C T C | A T G | C C C | T T C | G G C | T G C | C T C | C T G | G A C |
| **PRKCA KO Osimertinib #1** | C T C | A T C | A C G | C A G | C T C | A T G | C C C | T T C | G G C | T G C | C T C | C T G | G A C |
| **PRKCA KO Osimertinib #6** | C T C | A T C | A C G | C A G | C T C | A T G | C C C | T T C | G G C | T G C | C T C | C T G | G A C |
| **PRKCA KO Osimertinib #7** | C T C | A T C | A C G | C A G | C T C | A T G | C C C | T T C | G G C | T G C | C T C | C T G | G A C |
| **PRKCA KO Osimertinib #8** | C T C | A T C | A C G | C A G | C T C | A T G | C C C | T T C | G G C | T G C | C T C | C T G | G A C |

***Note:*** *Key sites associated with erlotinib or osimertinib resistance* (*T790M and C797X respectively) are highlighted. Only available endpoint tumors were sequenced.*

**Table S2** TaqMan and siRNA Sequences/Identifiers

| Target | Sequence | Source | Assay ID | Assay |
|---|---|---|---|---|
| **siPRKCA** | CAACGUACCCAUUCCGGAATT | Thermo Fisher | Cat#S11092 | siRNA |
| **siALDH1A1** | AAGGATTACCCTTCCAACGAA | Qiagen LLC | Cat#SI00011697 | siRNA |
| **All-Star** | Scramble Control siRNA (siAS) | Qiagen LLC | Cat#1027281 | siRNA |
| **ALDH1A1** | | Thermo Fisher | Hs00946916_m1 | TaqMan |
| **CD24** | | Thermo Fisher | Hs00897386_m1 | TaqMan |
| **CD36** | | Thermo Fisher | Hs00354519_m1 | TaqMan |
| **CD70** | | Thermo Fisher | Hs00174297_m1 | TaqMan |
| **CD133** | | Thermo Fisher | Hs01009259_m1 | TaqMan |
| **CPT1A** | | Thermo Fisher | Hs00912671_m1 | TaqMan |
| **CPT1C** | | Thermo Fisher | Hs00380581_m1 | TaqMan |
| **GPX4** | | Thermo Fisher | Hs00989766_g1 | TaqMan |
| **KDM5B** | | Thermo Fisher | Hs00231908_m1 | TaqMan |
| **NFE2L2** | | Thermo Fisher | Hs00975961_g1 | TaqMan |
| **PRKCA** | | Thermo Fisher | Hs00925195_m1 | TaqMan |
| **PRKCB** | | Thermo Fisher | Hs00176998_m1 | TaqMan |
| **PRKCG** | | Thermo Fisher | Hs00177010_m1 | TaqMan |
| **PRKCD** | | Thermo Fisher | Hs01090047_m1 | TaqMan |
| **PRKCE** | | Thermo Fisher | Hs00942886_m1 | TaqMan |
| **PRKCH** | | Thermo Fisher | Hs00178933_m1 | TaqMan |
| **PRKCQ** | | Thermo Fisher | Hs00234709_m1 | TaqMan |
| **PRKCI** | | Thermo Fisher | Hs00995852_g1 | TaqMan |
| **VIM** | | Thermo Fisher | Hs00185584_m1 | TaqMan |
| **ACTB** | | Thermo Fisher | Hs01060665_g1 | TaqMan |

**Supplemental References**


1. Cancer Cell Line Encyclopedia (RRID:SCR_013836).
